# Direct induction of hemogenic endothelial progenitors from hPSCs by defined factors revealed by single-cell transcriptome analysis

**DOI:** 10.1101/2021.03.18.435636

**Authors:** Lauren N. Randolph, Yuqian Jiang, Yun Chang, Xiaoping Bao, Xiaojun Lance Lian

**Affiliations:** Department of Biomedical Engineering, Pennsylvania State University, University Park, PA, 16802, USA; Department of Biology, Pennsylvania State University, University Park, PA, 16802, USA; The Huck Institutes of the Life Sciences, Pennsylvania State University, University Park, PA, 16802, USA; Davidson School of Chemical Engineering, Purdue University, West Lafayette, IN, 47907, USA

**Keywords:** human pluripotent stem cells, hematopoietic differentiation, SOX17, forward programming

## Abstract

Transcription factors (TFs) play critical roles in stem cell maintenance and differentiation. Using single cell RNA sequencing, we investigated TFs expressed in hemogenic endothelial (HE) progenitors differentiated from human pluripotent stem cells (hPSCs) and identified upregulated expression of SOXF factors *SOX7, SOX17*, and *SOX18* in the HE population. To test whether overexpression of these factors increases HE differentiation efficiency, we established inducible hPSC lines and found only *SOX17* improved differentiation. Temporal expression analysis further revealed *SOX17* was turned on immediately before VE-Cadherin, indicating SOX17 may be a causative factor for HE differentiation. Upon *SOX17* knockdown via CRISPR-Cas13d, HE differentiation was significantly abrogated. Strikingly, we discovered *SOX17* overexpression alone is sufficient to generate more than 50% CD34^+^VE-cadherin^+^CD73^-^ cells that could be directed to hematopoietic progenitors, which emerged via an endothelial-to-hematopoietic transition and significantly upregulated definitive hematopoietic transcriptional programs. Functional assays showed that these progenitors can differentiate into blood cells from multiple lineages. Our analyses reveal an uncharacterized function of *SOX17* in directing hPSCs differentiation towards HE cells.

**Significance Statement:** Hemogenic endothelial (HE) cells have been generated from human pluripotent stem cells (hPSCs) to study blood development. However, their full transcriptomic characterization and key genes involving in directing HE differentiation is unclear. Utilizing single cell RNA-seq analysis, we find that SOX17 is solely expressed in HE cells and is also required for HE differentiation. Strikingly, we find that overexpression of SOX17 alone is sufficient to program hPSCs into CD34+VE-cadherin+CD73-HE cells, which could further differentiate into blood progenitors. Our research reveals that SOX17 is sufficient to direct hPSCs differentiation to HE cells.

**Classification:** Physical Sciences/Engineering; Biological Sciences/Cell Biology.

## Main Text

### Introduction

Human pluripotent stem cell (hPSC) differentiation via growth factors and small molecules often results in heterogenous populations of cells, dramatically affecting our ability to efficiently derive therapeutically relevant cell types (1–4). Generally speaking, a somatic cell, which emerges late in human development, is more difficult to derive because the mechanistic basis of that cell’s development may be largely unknown. For example, hematopoietic development occurs at several developmental timepoints thereby increasing the complexity. Primitive hematopoiesis is defined as occurring in the yolk sac and does not give rise to lymphoid cells or hematopoietic stem cells (HSCs) (5). A second hematopoietic event localized in the yolk sack produces erythroid-myeloid progenitors as well as lymphoid progenitor cells (5). Despite closer resemblance to adult blood components, this hematopoietic event does not produce definitive HSCs (5). Chronologically the third stage of hematopoiesis in mammals is the first to generate HSCs capable of long-term, multi-lineage blood reconstitution and is thus termed definitive hematopoiesis (6). Definitive hematopoietic cells develop from hemogenic endothelial (HE) progenitors via an endothelial-to-hematopoietic transition (EHT) process (5–7). Due to the lack of complete understanding of molecular pathways controlling definitive HSC development, it is challenging to efficiently derive HE progenitors and HSCs from hPSCs.

An alternative method to derive desired cell types from hPSCs is forward programming, which is defined by overexpression of appropriate TFs in hPSCs to program them into desired cell types. Forward programming can generate desired cells with a high efficiency in a much shorter period of time. For example, *NFIA* overexpression is sufficient to generate astrocytes from neural stem cells within 3 weeks, as compared to 3-6 months using growth factor/small molecule-based protocols (8). To apply forward programming to deriving HE cells, it is critical to know which TFs are specifically expressed in this population.

With the development of single cell analysis techniques, researchers are able to characterize differentiated cell types at a depth previously unattainable. Notably, single-cell RNA sequencing (scRNA-seq) analysis reported heterogeneity in many differentiated populations, including endothelial cells and pancreatic beta cells (9, 10). It follows that scRNA-seq can be used to identify critical TFs solely expressed in desired cell populations, which may be used for forward programming. Using scRNA-seq analysis, we reveal that SOXF factors (11), *SOX17, SOX7*, and *SOX18*, are specifically expressed in our hPSC derived HE progenitors. We systematically study *SOX17* during hPSC differentiation to HE progenitors and illustrate that *SOX17* overexpression enhances HE differentiation. Knockdown of *SOX17* via CRISPR-Cas13d, however, inhibited HE progenitor differentiation. Importantly, overexpression of SOX17 alone in the absence of any small molecules or growth factors is sufficient to differentiate hPSCs into HE progenitors that can further differentiate into multiple hematopoietic lineages. In summary, our findings provide new insights into HE development and point out a critical role of SOX17 in differentiating hPSCs into HE cells, enabling a novel forward programming method to generate HE cells via overexpression of SOX17 in hPSC.

## Results

### Single cell RNA sequencing reveals SOXF factors expression in hemogenic endothelial progenitors

We previously developed a protocol to generate endothelial progenitors using an initial pulse of Wnt/β-catenin signaling activation with a GSK3β inhibitor, CHIR99021 (CH) (1, 12, 13). These progenitors express CD34, CD31, and VE-Cadherin (VEC), generate primitive vascular structures, and have recently been shown to give rise to hematopoietic cells (1, 14). To further establish if our differentiation protocol results in HE cells and identify TFs enriched in the HE population, we performed single cell transcriptome analysis of day 5 differentiated cells (**Fig. 1A**). We sequenced 2,673 cells and after quality control filters were applied, we included 1,917 cells in our analysis (**Fig. S1A**). Dimensional reduction and supervised clustering showed five distinct clusters of cells on day 5 of differentiation (**Fig. 1B-C, S1B-D**). Clusters (0, 1, 2, 3, and 4) were composed of 609, 457, 454, 358, and 39 cells respectively. Based on the top 100 differentially expressed genes, cluster 0 likely represents cardiac progenitors with upregulation of *TNNI1, HAND1*, and *TMEM88* (15) (**Fig. 1C, Table S1**). Cluster 3 cells show differential expression of *MYL7, MYL9*, and *HAND1*, and may be therefore labeled as atrial cardiac progenitors (**Table S1**). Cluster 4 has increased expression of *SOX2 and POU5F1* and may represent residual undifferentiated hPSCs (**Table S1**). Cluster 1 separated further away from the other four clusters in the UMAP projection and showed increased expression of *CD34, CDH5* (*VEC*), and *PECAM1* (*CD31*), indicating their endothelial progenitor identity. Other genes that have been identified as markers of the HE population or early hematopoietic lineages, including *CD93, MECOM, ETS1, KDR, KIT*, and *RUNX1*, showed increased expression in cluster 1 as compared with all other clusters (**Fig. S1E-J**) suggesting their HE identity (3, 16–20). While increased resolution further separated cluster 1 into two clusters, we did not observe distinct segregation of HE markers or type of endothelial cells (**Fig. S1K**). Both groups showed increased expression of DLL4 and NOTCH1, indicative of arterial type endothelial cells, as well as CD34 expression, although CD34 was expressed at higher levels in one of the two sub-clusters. (**Fig. S1L-N**). Venous endothelial gene expression was also observed in both clusters although was higher in cluster 5 (**Fig. SO-P**).

**Figure 1.**
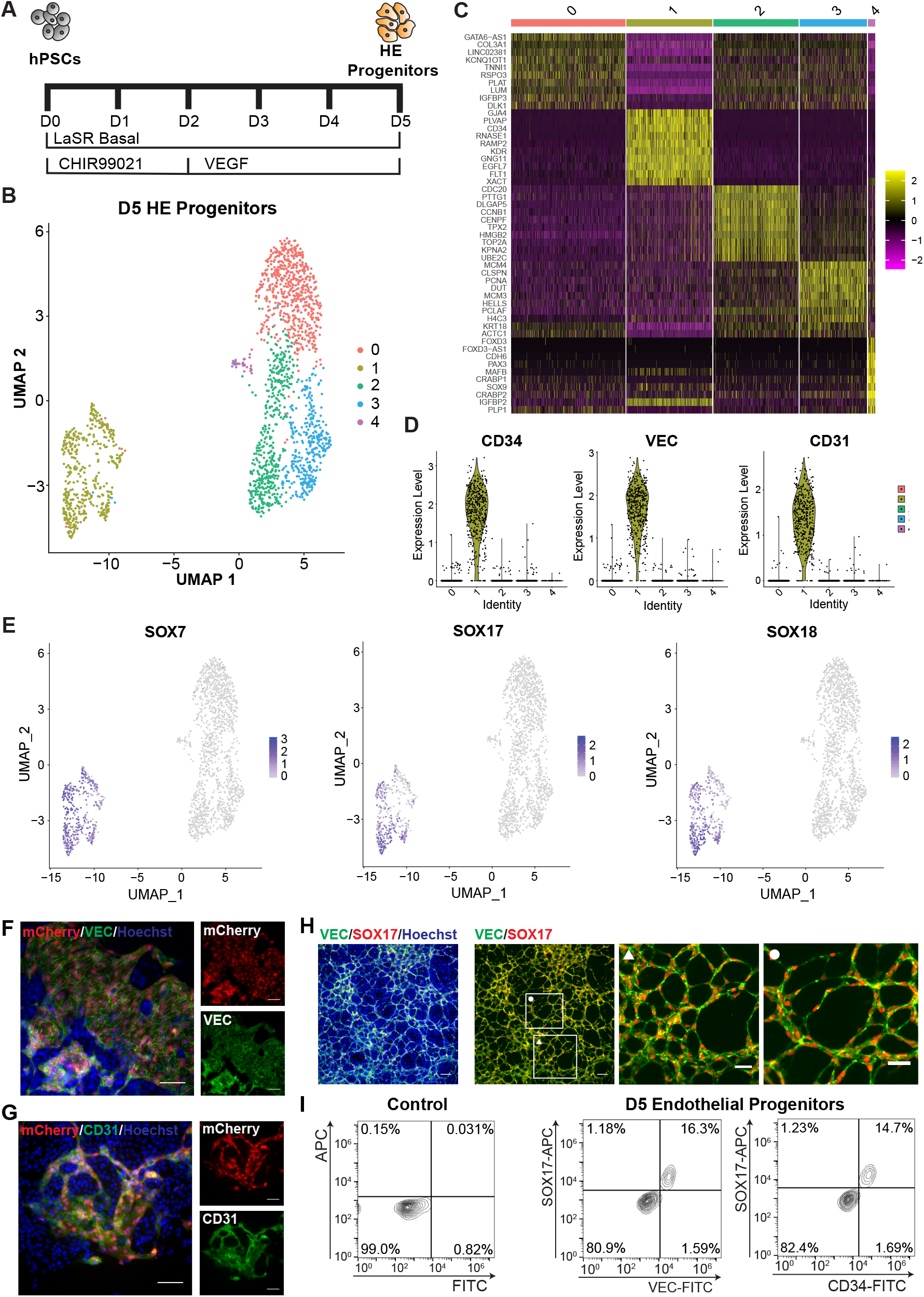
CH-induced endothelial progenitor differentiation method yields population expressing hemogenic endothelial markers and SOXF family. **A)** Schematic of endothelial progenitor differentiation used to test for hemogenic potential. **B)** UMAP dimensional reduction projection showing clustering of cells produced on D5. **C)** Heat map indicating the top 10 variably expressed genes for each cluster. **D)** Violin plots identifying CD34, VEC, and CD31 expression in cluster 1. **E)** SOXF family member expression in D5 population. **F-G)** Immunofluorescence images of day 5 endothelial progenitors derived from SOX17-mCherry reporter cells. mCherry expression is seen in cells that express VEC **(F)** and CD31 **(G)**. Nuclear staining was done with Hoechst 33342. Scale bars are 100 µm. **H)** Immunofluorescent images showing VEC and SOX17 co-expression in day 5 cells differentiated from 6-9-9 cells. White boxes indicate locations of enlarged views with symbols indicating the corresponding image. Nuclear staining was done with Hoechst 33342. Left two scale bars are 100 µm and right two scale bars are 50 µm. **I)** Flow cytometry analysis showing co-expression of SOX17 and VEC and SOX17 and CD34 in day 5 cells.

Inspection of differentially expressed TFs uncovered high expression of all three SOXF factors in our HE cells (**Fig. 1E**). No other SOX factors were differentially expressed in cluster 1; although we did see differential expression of SOX6 and SOX4 in cluster 0 and SOX2 and SOX9 in cluster 4 (**Table S1**). To confirm SOXF expression in HE cells, we used a SOX17-mCherry knockin reporter hPSC line (21) and differentiated the knockin hPSCs to day 5 differentiated cells. The reporter cell line was validated by antibody staining on day 5 of differentiation (**Fig. S1Q**). In addition, mCherry expression only occurred in cells also expressing VEC and CD31 on day 5 of differentiation (**Fig. 1F-G**). The co-expression of SOX17 with HE progenitor markers VEC and CD34 was also observed in the HE differentiation of an additional hPSC line (6-9-9) by immunostaining and flow cytometry (**Fig. 1H-I**). Furthermore, SOX17 in particular has been identified as a marker of definitive HE cells in both mouse and human models (22–24). These data, along with a study (14) in which Galat *et al*. generated lymphoid cells from endothelial progenitors using our CH-induced differentiation protocol, provide strong evidence that our protocol does result in HE cells and our scRNA-seq data revealed upregulated expression of SOXF factors in HE cells.

### Overexpression of SOX17 but not SOX7 or SOX18 enhances hemogenic endothelial differentiation efficiency from hPSCs

Upon discovering all 3 SOXF factors are expressed in HE cells, we sought to understand which, if any of these factors, play a functional role in determining HE cell fate. To address this question, we generated cell lines with inducible overexpression of *SOX7, SOX17*, and *SOX18* by cloning each TF into our doxycycline (Dox) inducible, PiggyBac-based XLone construct (25) (**Fig. 2A**). This construct was then introduced into hPSCs, and cells successfully incorporating the construct were purified by drug selection (**Fig. 2A**). The modified cells were then referred to as H9+XLone-SOX7, H9+Xlone-SOX17, and H9+XLone-SOX18. To ensure the desired function of resulting cells, H9+XLone-SOX17 cells were treated with or without Dox for 24 hours, which revealed robust overexpression of SOX17 up to 82.6%, with minimal leakage in cells not treated with Dox (**Fig. 2B**). This was consistent with our previous findings and indicates the successful generation of stable transgenic cell lines (25).

**Figure 2.**
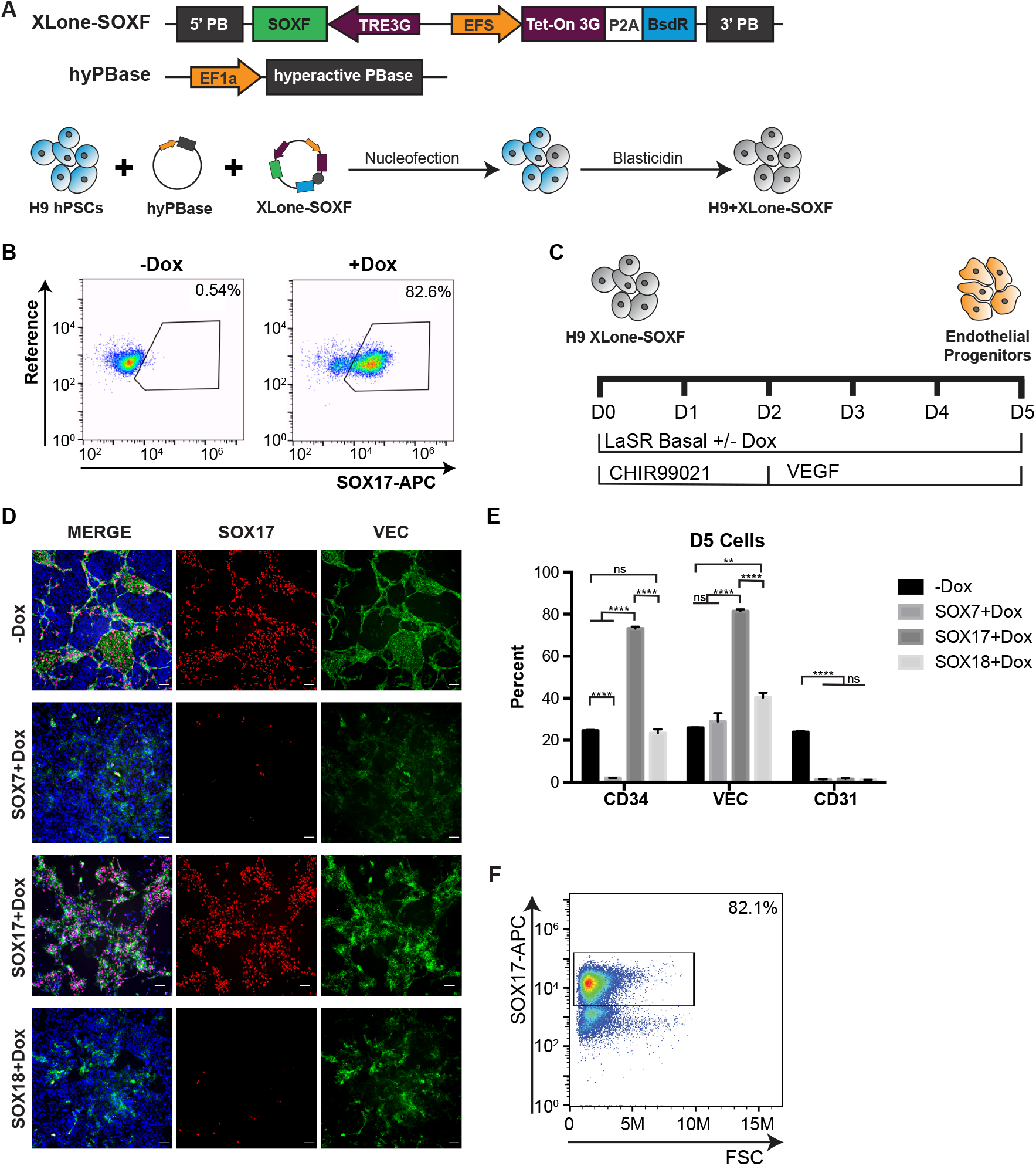
SOX17 is the only SOXF member that increases HE progenitor differentiation efficiency when overexpressed. **A)** Schematic showing the generation of H9+XLone-SOXF (SOX7, SOX17, or SOX18) hPSCs. **B)** Flow cytometry analysis of SOX17 expression in H9+XLone-SOX17 cells treated with or without Dox for 24 hours. **C)** Schematic illustrating culture conditions for SOXF overexpression during HE progenitor differentiation. **D)** Immunofluorescence analysis of day 5 cells differentiated with Dox for each XLone-SOXF cell line and positive control without Dox. Cells were stained with SOX17 and VEC antibodies and nuclear stain (Hoechst 33342). Scale bars are 100 µm. **E)** Quantification of flow cytometry analysis of CD34, VEC, and CD31 expression in D5 H9+XLone-SOXF cells treated with Dox and control without Dox (n=3). ** and **** indicate p < 0.01 and p < 0.0001 respectively. Error bars represent standard error of the mean. **F)** Flow cytometry analysis of SOX17 expression in D5 H9+XLone_SOX17 cells differentiated with Dox.

To test whether overexpression of SOXF factors enhances CH-induced HE differentiation from hPSCs, we differentiated each cell line to HE progenitors in the presence or absence of Dox and compared the expression of CD34, VEC, CD31, and SOX17 (**Fig. 2C, S2A**). The expression levels of these markers in cells without Dox treatment was consistent with our earlier experiments, and the differentiation efficiency was comparable with our previous results (1, 13) (**Fig. 2D-E, S2B**). For cells differentiated in the presence of Dox, we observed a dramatic increase in the SOX17^+^ population of up to 82% for the H9+XLone-SOX17 cells, as would be expected, and very few SOX17^+^ cells in the day 5 cells generated from either H9+XLoneSOX7 or H9+XLone-SOX18 (**Fig. 2D and F**). H9+XLone-SOX7 cells also significantly decreased the percentage of CD34+ cells with no change in the percentage of VEC^+^ cells (**Fig. 2D-E, S2B**). H9+XLone-SOX18 cells showed no change in the CD34^+^ population but a statistically significant increase in the VEC^+^ population (40.27% ± 2.32%, p = 0.0035) (**Fig. 2D-E, S2B**). The day 5 cells resulting from H9+XLone-SOX17 cells showed significant increases in CD34^+^ (73.10% ± 0.94%, p = 1.15e-6) and VEC^+^ (81.40% ± 0.90%, p = 4.97e-7) populations over differentiation without any transgene expression (**Fig. 2D-E, Supplementary Fig. 5**). Fluorescent microscopy revealed the majority of SOX17^+^ cells were also VEC^+^ (**Fig. 2D**). We did not observe a statistically significant difference in the yields for cells generated with each SOXF factor; however, there were fewer cells generated than with the normal differentiation protocol (**Fig. S2C**). Interestingly, overexpression of each SOXF factor resulted in a loss of CD31 expression (**Fig. 2E, S2B**), indicating that SOXF overexpression may convert hPSCs directly to CD34+VEC+CD31-stage cells. Collectively, this demonstrates that forced overexpression of *SOX17*, but not *SOX7* or *SOX18*, increases CH-induced HE differentiation efficiency, highlighting a functional role for *SOX17* in the acquisition of HE cell fate.

### SOX17 expression occurs prior to hemogenic endothelial markers and CRISPR-Cas13d interference with SOX17 inhibits HE differentiation

To better understand the role of SOX17 during differentiation, we characterized SOX17 expression kinetics along differentiation. We differentiated hPSCs following the protocol illustrated in **Fig. 1A** and collected cells daily until day 3 and every six hours after day 3 until day 5. Western blot analysis showed SOX17 and VEC are first detected on day 3.75 (**Fig. 3A-B, S3A**). Immunofluorescent analysis revealed there are more cells expressing SOX17 alone on day 3.75, and this SOX17 single positive cell number gradually decreases as cells become double positive for SOX17 and VEC over time (**Fig. 3C-D, S3B**). This data indicates expression of SOX17 occurred before the appearance of VEC^+^ HE cells.

**Figure 3.**
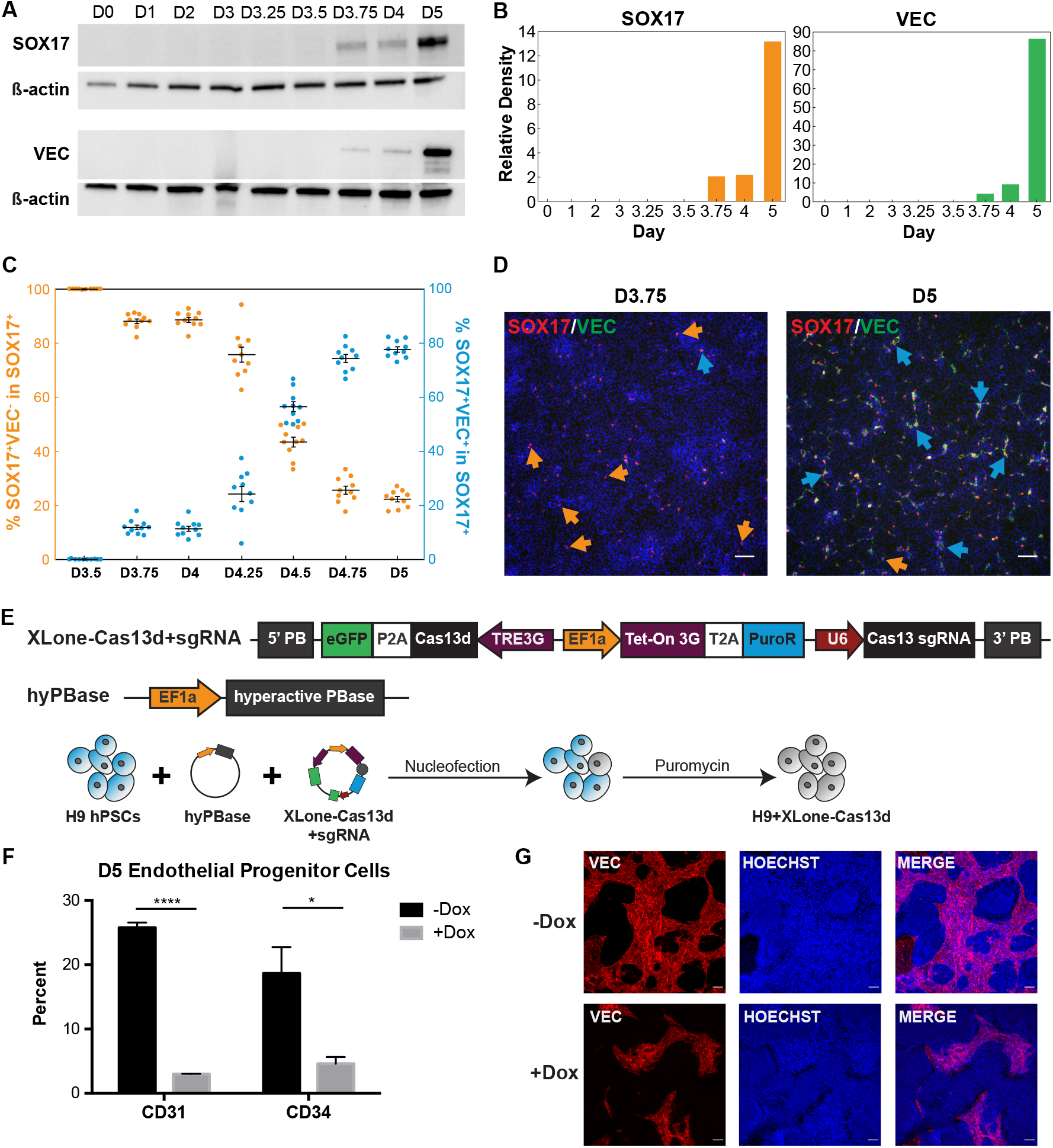
SOX17 expression occurs immediately prior to endothelial marker expression and transcriptional interference reduces endothelial marker expression. **A)** Western blots showing SOX17 and VEC protein levels over the course of differentiation. ß-actin was used as a housekeeping gene. **B)** Quantification of blots shown in **(A)** normalized to ß-actin. **C)** Quantification of SOX17^+^VEC^-^ cells (left axis, orange) and SOX17^+^VEC^+^ cells (right axis, blue) in the SOX17^+^ population as derived from analysis of immunofluorescent images (n=10). Error bars represent standard error of the mean. **D)** Representative immunofluorescence images used for quantification in **(C)** for day 3.75 and day 5 (n=10). Orange arrows highlight SOX17^+^VEC^-^ cells and blue arrows highlight SOX17^+^VEC^+^ cells. Nuclear staining was done with Hoechst 33342. Scale bars are 100 µm. **E)** Schematic illustrating the generation of a cell line with Cas13d-based inducible SOX17 knockdown. **F)** Quantification of flow cytometry experiments analyzing the change in size of the day 5 population expressing CD31 or CD34 for cells treated with and without Dox (n=3). * indicates p < 0.05 and **** indicates p < 0.0001. Error bars represent standard error of the mean. **G)** Immunofluorescence images of D5 cells differentiated with or without Dox stained with VEC. Nuclear staining was done with Hoechst 33342. Scale bars are 100 µm.

To further study the role of SOX17 in HE differentiation, we sought to perform a loss-of-function analysis using a CRISPR-Cas13d-meditated knockdown approach. This member of the Cas13 family can knockdown RNA transcripts efficiently (26). We cloned Cas13d into our XLone plasmid construct under the control of the inducible TRE3G promoter (**Fig. 3E**) (25). We also cloned a U6 promoter expressing SOX17 gRNA into the plasmid to establish a single transposon system for Cas13d interference. We nucleofected this plasmid into H9 cells and used puromycin drug selection to purify the cells that integrated XLone-Cas13d system (**Fig. 3E**). To test Cas13d-mediated SOX17 knockdown efficiency, definitive endoderm (SOX17+ cells) differentiation with or without Dox was performed and this experiment revealed robust and near complete SOX17 knockdown was achieved as measured by flow cytometry analysis (**Fig. S3C**). HE differentiation of these cells with and without Dox treatment revealed abrogation of CD31 and CD34 expression as well as reduced VEC expression upon SOX17 knockdown (**Fig. 3F-G, S3D**). These results demonstrate that SOX17 expression is required for the formation of HE cells from hPSCs.

### Overexpression of SOX17 alone is sufficient for the generation of cells expressing CD34 and VEC from hPSCs

To evaluate the effects of *SOX17* overexpression alone on the differentiation of hPSCs, we treated transgenic XLone-SOX17 hPSCs with or without Dox in a basal medium (**Fig. S4**). A loss of pluripotency was observed by a notable loss of stem cell morphology as early as day 1 (**Fig. S4**). Day 5 cells cultured in basal media without Dox did not turn on expression of CD34, CD31, or VEC; however, when Dox treatment is applied, a population of the cells show expression of CD34 and VEC, with CD34 RNA expression increasing as early as day 1 (**Fig. 4A-C, S5**).

**Figure 4.**
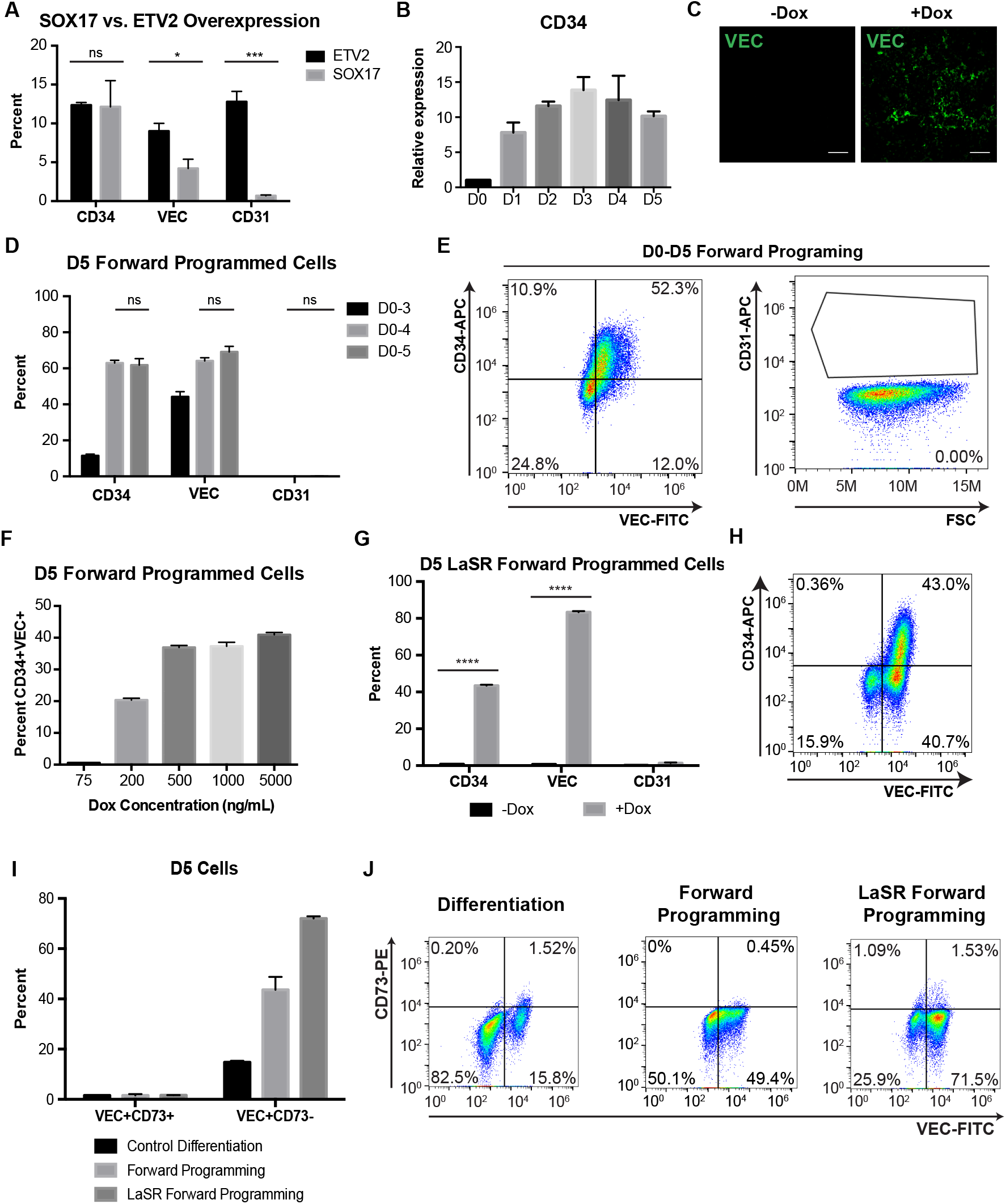
SOX17 forward programming is sufficient to produce CD34^+^VEC^+^ cells. **A)** Quantification of flow cytometry analysis of day 5 cells for endothelial progenitor markers VEC, CD34, and CD31. Day 5 SOX17 forward programmed cells are compared to ETV2 forward programmed cells. *** indicates p < 0.001. **B)** qPCR analysis of CD34 expression from day 0 to 5 for LaSR Basal +Dox condition. **C)** Immunofluorescence analysis of VEC expression in day 5 cells for each condition. Scale bars are 130 µm. **D)** Quantification of flow cytometry analysis for day 5 cells for endothelial progenitor markers CD34, VEC, and CD31 for cells treated with Dox for 3, 4, and 5 days. **E)** Representative flow cytometry plots showing SOX17 forward programmed cells with Dox treatment for 5 days. **F)** Quantification of flow cytometry analysis for different Dox concentrations used for 5 days of treatment. **G)** Quantification of day 5 flow cytometry data for day 5 cells cultured in LaSR hPSC media with or without Dox. **** indicates p < 0.0001. **H)** Representative flow cytometry analysis of day 5 cells treated with or without Dox in LaSR hPSC media. **I)** Quantification of flow cytometry analysis of CD73 and VEC expression in day 5 cells. **J)** Representative flow cytometry plots for CD73 and VEC flow analysis.

In view of the fact that other TFs such as *ETV2* have been used to forward program hPSCs to endothelial progenitors (27), we compared marker expression for the cells resulting from *ETV2* and *SOX17* forward programming. We used the same XLone plasmid construct to establish an H9+XLone-ETV2 cell line (**Fig. S5A-D**). These cells were then forward programmed in basal media with or without dox and analyzed on day 5. Compared to *SOX17* forward programmed cells, there was no significant difference in CD34 (p = 0.95); however, there was a significant increase in VEC (p = 0.037) and CD31 (p = 0.0008) expression with *ETV2* (**Fig. 4A**). The difference in marker profile could be indicative of a difference in progenitor potential. To compare further endothelial cell potential, the day 5 cells were further differentiated to day 20 using commercial endothelial cell media. After 20 days, the *ETV2* forward programmed progenitors resulted in cells that strongly expressed mature endothelial markers CD31, VEC, vWF, and I-CAM1 (**Fig. S5F**). These markers were not detected in *SOX17* forward programmed cells, indicating these cells did not further differentiate into endothelial cells via endothelial cell media treatment (**Fig. S5E**).

To confirm that SOX17-mediated forward programming was not cell line dependent, we generated and validated XLone-SOX17 cell lines using another hPSC line (H1 OCT4-GFP cells) (**Fig. S6A-B**). Cells were then cultured with or without Dox treatment in basal media (**Fig. S6C**). We confirmed the loss of pluripotency by decrease in the GFP expression for H1 OCT4-GFP+XLone-SOX17 cells from 97.1% on day 0 to 20.8% on day 5 (**Fig. S6D**).

To further increase the efficiency of *SOX17* forward programming, we hypothesized that duration of Dox treatment and/or passaging cells during differentiation may play a role in increasing efficiency. We forward programmed hPSCs, passaging them on day 2, and varied the duration of Dox treatment from 3 to 5 days (**Fig. 4D**). With day 2 passaging, we found the CD34^+^ population generated from 3-day Dox treatment was similar to that of forward programming without a day 2 passage (**Fig. 4A and D, S7A**). However, the size of the VEC^+^ population increased from less than 5% (no day 2 passaging) to 44.2% ± 2.8% (with day 2 passaging) (**Fig. 4A and D, S7A**). Longer duration of Dox treatment increased the efficiency to at least 51% ± 2.3% CD34+VEC+ with no significant difference between 4 or 5 days of Dox treatment (CD34 p = 0.79, VEC p = 0.23) (**Fig. 4D-E, S7A**). This demonstrated that longer Dox treatment and passaging on day 2 increased the efficiency of *SOX17* forward programming. The passage could serve as selection for cells that are being forward programmed or the change in cell-cell contact could further enhance differentiation.

After optimizing the required duration of Dox treatment, we tested a range of concentrations to determine the optimal level of transgene activation (25). We found that lower Dox concentrations resulted in a decreased CD34^+^VEC^+^ population and that at least 500 ng/mL Dox was required to achieve maximal efficiency (**Fig. 4F, S7B**). This indicates that maximal transgene expression is required for *SOX17* forward programming. We replicated this finding with another hPSC line (6-9-9+XLone-SOX17) obtained 57.2% ± 2.8% of the resulting cells expressing CD34 and VEC (**Fig. S7C**).

In an effort to examine the potency of SOX17 as a mediator of forward programming, we tested the effects of *SOX17* forward programming in hPSCs cultured in hPSC media. Cells were cultured in one of two hPSC media, LaSR or mTeSR1, in the presence or absence of Dox (**Fig. S8A-B**). *SOX17* forward programming was able to overcome pluripotency maintenance signals in hPSC media and cause cells to differentiate as evidenced by loss of pluripotent morphology (**Fig. S8A-B**). Day 5 flow cytometry analysis showed the presence of CD34^+^VEC^+^ cells and CD34^-^VEC^+^ cells (**Fig. 4G-H, S8C-D**). We did not observe expression of CD31 in the day 5 cells, similar to the observations made in basal media (**Fig. 4G, S8D**). Both hPSC media produced CD34^+^VEC^+^ cells and CD34^-^VEC^+^ cells, albeit at different efficiencies (**Fig. 4H, S8C**). This could be due to the increased amount of bovine serum albumin in mTeSR1 as compared to LaSR media; nevertheless, both media produced statistically significant populations of differentiated cells expressing CD34 (LaSR p = 5.75e-8, mTeSR1 p = 2.88e-6, +dox vs -dox) and VEC (LaSR p = 6.78e-9, mTeSR1 p = 1.18e-6, +dox vs -dox) (**Fig. 4G, S8D**). Collectively, these results demonstrated *SOX17* forward programming potency can overcome the presence of pluripotency growth factors in hPSC media to efficiently produce CD34^+^VEC^+^ cells.

HE differentiations often result in heterogenous populations of cells, as is common with many *in vitro* methods (2, 3, 19). To further characterize the cells obtained from *SOX17* forward programming, we assessed the expression of an additional marker, CD73, which is not expressed in HE populations but is expressed in other non-HE endothelial cells (2). We compared day 5 cells resulting from CH-induced differentiation, forward programming in basal media, and forward programming in hPSC media (LaSR Forward Programming) and found that none of these methods produced more than an average of 1.62% of the cells expressing both VEC and CD73. This indicates that forward programming does not result in contamination with this previously identified non-HE endothelial cell population (**Fig. 4I-J**).

### SOX17 forward programming is dependent on β-catenin

To study interaction between SOX17 forward programming and CH-induced differentiation, we differentiated H9+XLone-SOX17 cells to day 5 with and without Dox while varying the presence of CH and VEGF. For conditions treated without Dox, cells that received no Wnt activation (CH) showed none of the HE markers (CD34, VEC, and CD31) (**Fig. 5A-C, S9A**). Cells differentiated using only Wnt activation showed minimal CD34 CD31, or VEC (**Fig. 5A-C, S9A**). This is supported by previous findings showing that female cell lines require the addition of VEGF to differentiate to endothelial progenitors (13). Cells differentiated with Wnt activation and VEGF expressed CD31, CD34, and VEC (**Fig. 5A-C, S9A**). For conditions treated with Dox, CD34 expression increased from at most 11% ±1.01% without Dox treatment to 67.6% ±1.50% with Dox treatment (**Fig. 5B-C, S9B**). The VEC^+^ population increased with Dox treatment similar to our observations for CD34 (**Fig. 5A, S9B**). The addition of CH in the presence of *SOX17* overexpression did not significantly affect the percentage of the CD34^+^VEC^+^ population.

**Figure 5.**
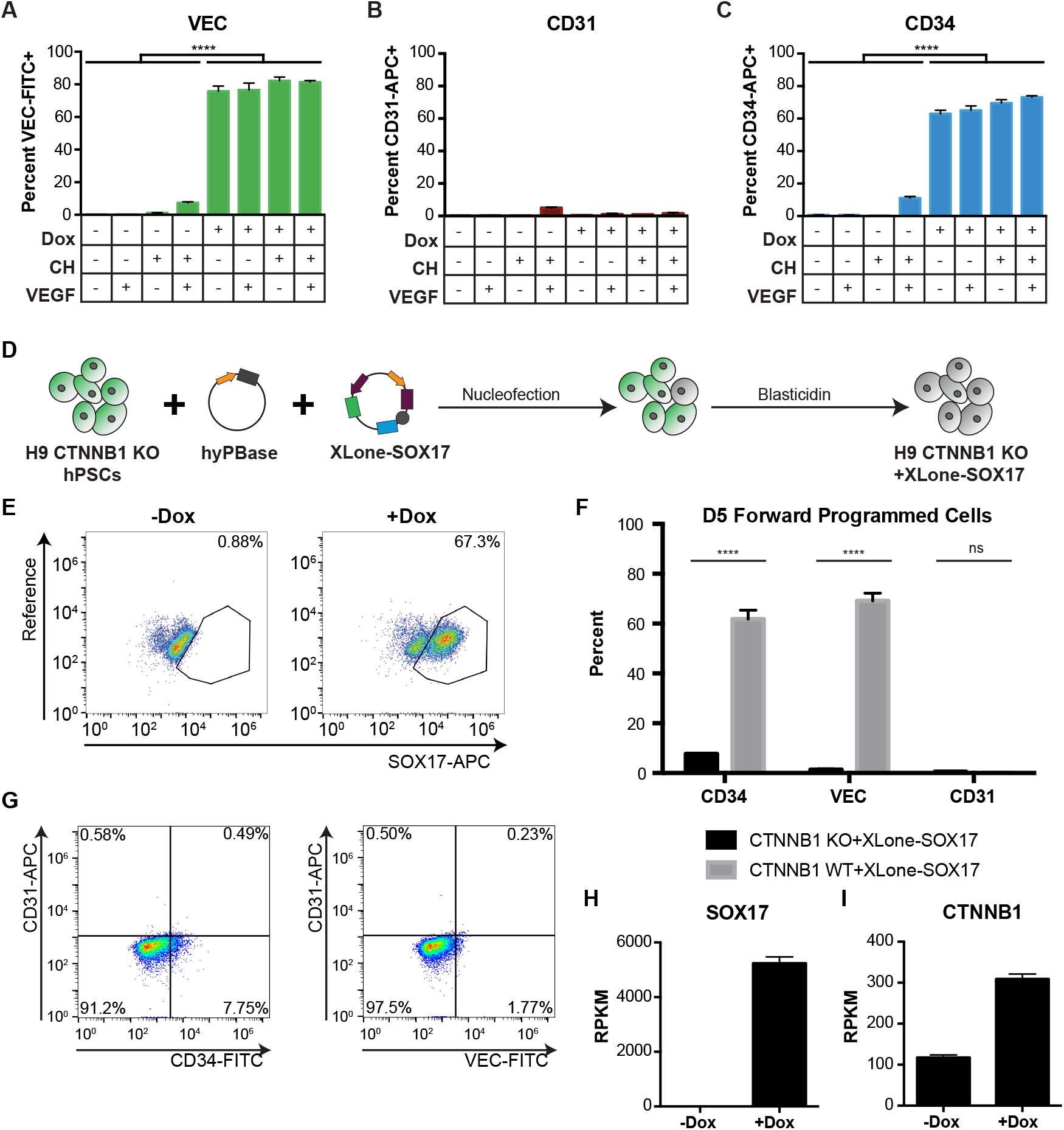
SOX17 forward programming is dependent on β-catenin. **A-C)** Quantification of flow analysis for endothelial progenitor markers VEC **(A)**, CD31 **(B)**, CD34 **(C)** with day 5 cells. **D)** Schematic showing the generation of H9 CTNNB1 KO +XLone-SOX17 hPSCs. **E)** Flow cytometry analysis validating the function of the H9 CTNNB1 KO +XLone-SOX17 cell line. **F)** Quantification of flow analysis for CD34, VEC, and CD31 for day 5 cells. **G)** Representative flow cytometry plots for H9 CTNNB1 KO +XLone-SOX17 day 5 forward programmed cells. **** indicates p < 0.0001. Error bars represent standard error of the mean and n=3 for all data. **H-I)** Inducible SOX17 hPSCs were used. Dox upregulated SOX17 expression **(H)** and CTNNB1 expression also upregulated **(I)** as a result of SOX17 overexpression.

Wnt/β-catenin signaling plays critical roles in mesoderm and HE specification from hPSCs (3, 5, 19, 28). Because CH addition did not further increase efficiency of *SOX17* forward programming, we sought to understand whether Wnt/β-catenin signaling is required for *SOX17* forward programming. To do this, we examined the efficacy of forward programming in the absence of β-catenin. We used our established H9 CTNNB1 KO cells (29), introduced the XLone-SOX17 construct, and verified inducible cell line function (**Fig. 5D-E**). These cells were then forward programmed with Dox in basal media for five days. The resulting day 5 cells showed a statistically significant decrease in the expression of CD34 (p = 5.52e-5) and VEC (p = 1.26e-5) as compared to cells with β-catenin (**Fig. 5F-G**). To examine whether CTNNB1 expression is upregulated when SOX17 is overexpressed, we analyzed RNA-seq data of Dox inducible SOX17 hPSCs with and without SOX17 overexpression (30). We confirmed that SOX17 was robustly overexpressed when treated with Dox and found that CTNNB1 was also upregulated as a result (**Fig. 5H-I**). This illustrated that β-catenin is required in *SOX17* forward programming.

### Further differentiated SOX17 forward programmed cells strongly upregulate hematopoietic transcription factors and yields cells capable of multilineage hematopoiesis

Based on our results showing SOX17 plays a critical role in HE progenitor acquisition, we sought to understand the extent to which SOX17 forward programming activates downstream hematopoietic gene programs. We forward programmed cells to day 5 via SOX17 overexpression and then removed Dox and cultured the cells in StemLine II hematopoietic stem cell expansion media (**Fig. 6A**). By day 10, many floating cells were observed, and we collected them to assess hematopoietic marker expression via flow cytometry. We found that around 20% to 60% of the suspension cells were positive for CD34, CD45, and CD44 (**Fig. 6B-E**). We then collected floating cells on days 8, 10 and 11 for qPCR analysis.

**Figure 6.**
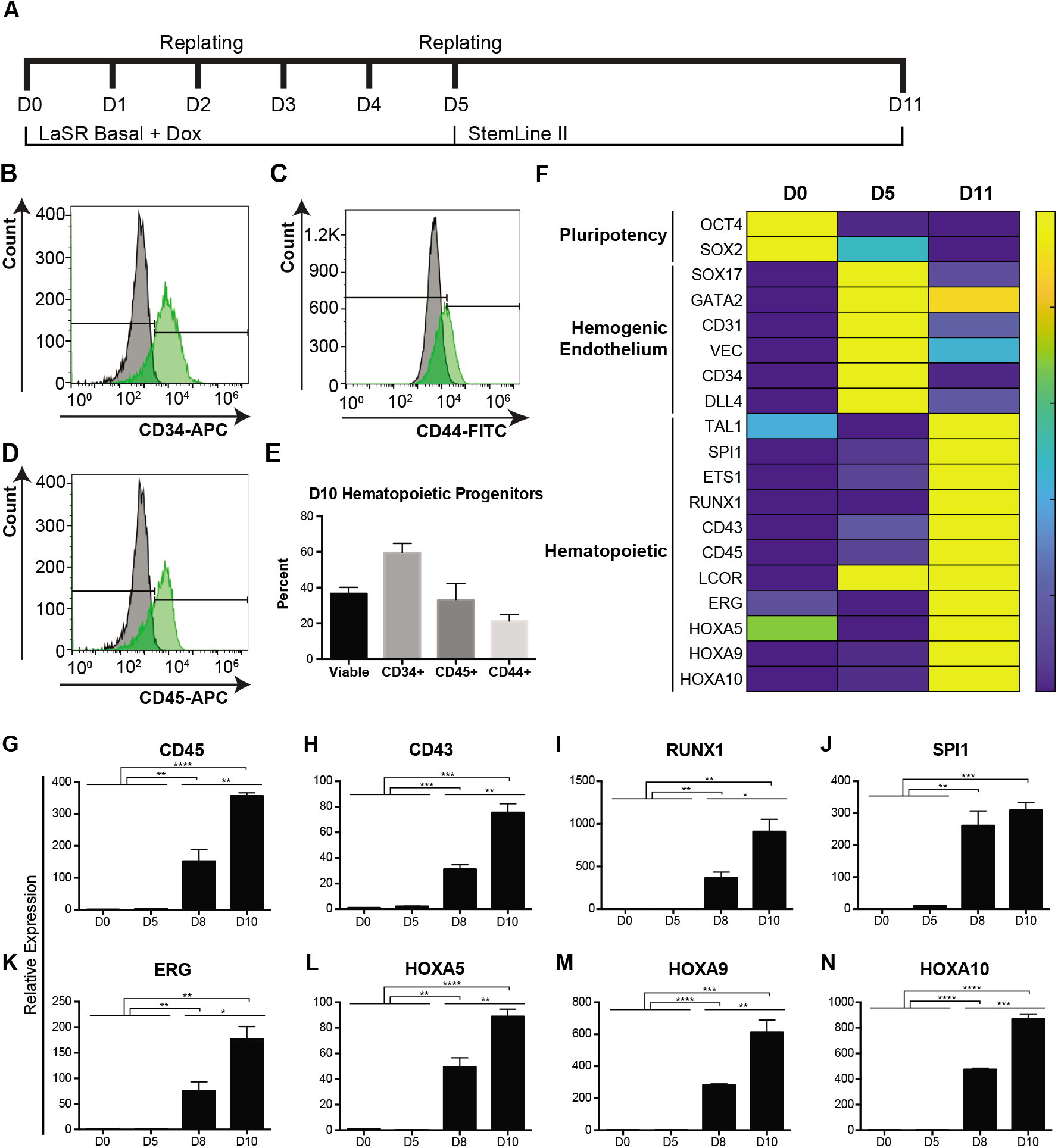
SOX17 forward programming results in hematopoietic progenitors that significantly upregulate hematopoietic transcription factors and surface markers. **A)** Schematic of defined and growth factor free hematopoietic progenitor differentiation with forward programming duration indicated by Dox treatment. **B)** Flow cytometry analysis of CD34 expression and cell viability for day 8 floating cells. **C)** Heat map showing qPCR data for day 0, day 5, and day 11 suspension cells (n=3). Data is depicted as expression relative to day 0 and scaled by row. **D-K)** Quantification of gene expression determined by qPCR (n=3) for CD45 **(D)**, CD43 **(E)**, RUNX1 **(F)**, SPI1 **(G)**, ERG **(H)**, HOXA5 **(I)**, HOXA9 **(J)**, and HOXA10 **(K)**. *, **, ***, and **** indicates p < 0.05, 0.01, 0.001, and 0.0001 respectively. Error bars represent standard error of the mean.

We performed qPCR analysis for genes associated with hPSCs, HE cells, and hematopoietic progenitors, including TFs previously used for hematopoietic reprogramming or forward programming. We observed stage specific peak expression for pluripotency genes (*OCT4* and *SOX2*) on day 0, HE genes (*SOX17, GATA2, VEC, CD34*, and *DLL4*) on day 5, and hematopoietic genes (*TAL1, SPI1, ETS1, RUNX1, CD43, CD45, LCOR, ERG, HOXA5, HOXA9*,

*HOXA10*) on day 11 (**Fig. 6F**). Further examination showed significant upregulation of hematopoietic surface markers *CD45* and *CD43* on both days 8 and 10, with 355-fold (p = 4.03e-6) and 75-fold (p = 0.0004) increases in relative expression respectively on day 10 (**Fig. 6G-H**). Furthermore, there was a significantly more CD45 expression in SOX17 forward programmed cells on day 10 as compared to control differentiated cells (p = 6.97e-5), SOX7 forward programmed cells (p = 4.28e-6), and SOX18 forward programmed cells (p = 4.31e-6) (**Fig. S10A**)

RUNX1 is a TF that plays a well-established role in hematopoietic fate acquisition and has been used for hematopoietic programming (31–33). We detected a significant increase in *RUNX1* on days 8 (p = 0.006) and 10 (p = 0.003) with expression approaching a 1000-fold increase over the day 0 expression level (**Fig. 6I**). Additionally, we observed strong upregulation of TFs *SPI1, ERG, HOXA5, HOXA9*, and *HOXA10* on days 8 and 10, with significant more expression than differentiated cells and SOX7 or SOX18 forward programmed cells (**Fig. 6J-N, S10B-C**). These factors along with RUNX1 have been used to program HE cells to a definitive hematopoietic fate (33). Taken together, these data demonstrate *SOX17* forward programming is sufficient for robust downstream activation of hematopoietic gene and TF networks.

To test the functional hematopoietic capacity of our SOX17 forward programmed hematopoietic progenitors, we performed a colony forming unit assay using floating cells obtained on days 8, 10, and 12. We observed all 5 types of colonies with the majority being erythrocytic (**Fig. 7A-B**). The presence of GM and GEMM colonies indicates that some of our progenitors are multipotent. We also performed a Giemsa stain on our day 10 floating cells to evaluate the types of cells already present after forward programming. We identified cells from all major blood lineages including cells with monocyte and lymphocyte morphology (**Fig. 7C**). This demonstrates the ability of our cells to differentiate to multiple hematopoietic cell lineages.

**Figure 7.**
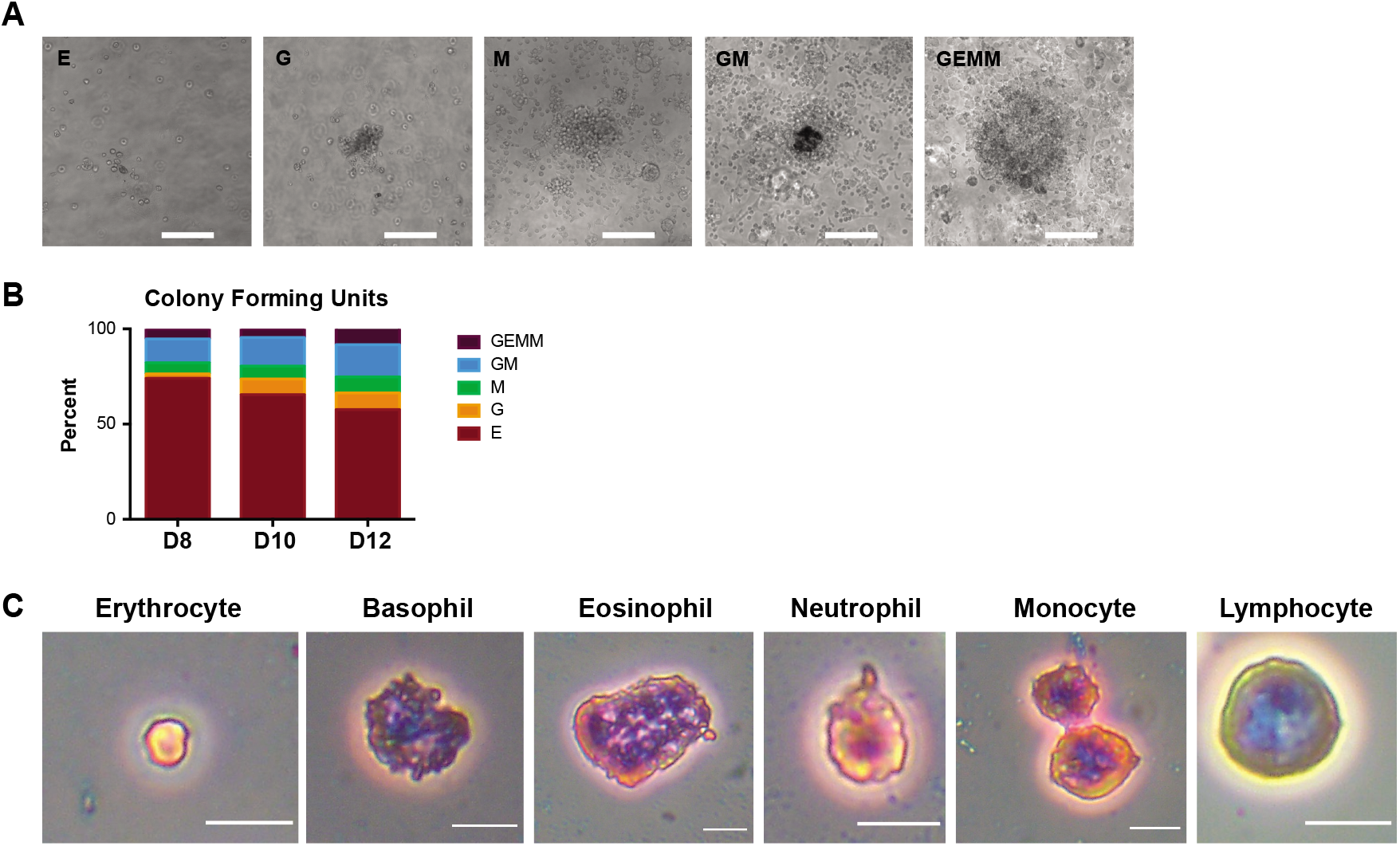
SOX17 forward programmed cells further differentiate to multiple hematopoietic lineages. **A)** Representative images of erythrocytic (E), granulocytic (G), macrophage (M), granulocytic/macrophage (GM), and granulocytic/erythrocytic/monocytic/megakaryocytic (GEMM) colony forming units. Scale bars are 200 µm. B) Quantification of colonies formed by day 8, 10, and 12 hematopoietic progenitors. C) Representative images of different hematopoietic cells obtained on day 10. Scale bars are 10 µm.

## Discussion

Forward programming has proven to be an effective strategy for the generation of difficult to obtain cell types(33). Herein, we have provided the first evidence to date that overexpression of *SOX17* alone in hPSCs is sufficient for the generation of CD34^+^VEC^+^CD73^-^ HE cells and floating CD34^+^ hematopoietic progenitors via an EHT in the absence of small molecules and growth factors. Furthermore, we demonstrated that the resulting hematopoietic progenitors have significantly upregulated key hematopoietic TFs and can further differentiate to multiple hematopoietic lineages. Single cell RNA sequencing of our published endothelial progenitor differentiation confirmed the expression of HE markers and revealed differential expression of SOXF factors. When these factors were overexpressed in tandem with differentiation, however, only SOX17 increased differentiation efficiency as measured by the generation of CD34^+^VEC^+^ cells. We did observe VEC expression with all three SOXF factors. This is consistent with reports that SOX17 can bind to the VEC promotor in human cells, and SOX7 binds to the VEC promotor and activates VEC expression in murine cells(34, 35). Given the high degree of conservation and the similarities in SOXF subgroup binding sites, it is likely all three factors will have a similar effect on the VEC transcription in human cells. This is supported by our findings indicating that SOX17 expression occurs just prior to VEC emergence. CD31 expression was not detected with *SOX17* forward programming; however due to the lack of HE specific markers that are not shared by other tissues and rarity of human HE cells during development it is difficult to determine definitively the marker profile of these cells. While the lack of CD31 expression was a departure from the marker profile we typically observe with our established CH-induced differentiation, other studies have characterized HE populations without the use of CD31 (19, 36). Furthermore, early progenitor populations have been identified that express high levels of CD34 and VEC and no CD31(37). It is also possible *SOX17* forward programming produces a different HE progenitor population lacking CD31 expression as is indicated by our comparison with *ETV2* forward programming.

To better understand the functional significance of SOX17, we performed loss of function studies using Cas13d and found SOX17 knockdown significantly impeded the ability of hPSCs to differentiate to HE cells. This is consistent with previous findings in murine and human systems(22, 34, 38). Having confirmed functional importance of SOX17 by loss of function, we went on to show that forward programming by *SOX17* overexpression is sufficient to obtain CD34^+^VEC^+^CD73^-^ cells from multiple hPSC lines in a variety of different culture media, including hPSC media. We optimized forward programming conditions to find the ideal temporal window and level for transgene expression. Furthermore, we found *SOX17* forward programming requires β-catenin indicating dependence on Wnt/β-catenin signaling, a hallmark of HE development(19). This is the first report to our knowledge that *SOX17* overexpression alone can efficiently generate HE cells.

Our data demonstrate overexpression of SOX17 during the first five days was sufficient to propel the cells through an EHT resulting in robust generation of CD34+ hematopoietic progenitors. Analysis of the expression of stage specific markers and TFs showed developmental progression from HE cells to hematopoietic progenitors. *SOX17* forward programmed cells were also able to strongly upregulate TFs associated with definitive hematopoietic fate and commitment. HOXA genes such as *HOXA5, HOXA9*, and *HOXA10* were significantly upregulated in hematopoietic progenitors that arose from *SOX17* forward programmed cells. This is consistent with our understanding that HOXA genes are only turned on by early definitive hematopoietic fate specification(7, 21). Furthermore, HOXA genes are selectively expressed in human fetal liver and umbilical cord blood derived hematopoietic progenitors(21).

Moreover, *SOX17* forward programming led to significant increases in expression for all of the TFs used by Sugimura *et al*. to forward program HE cells to definitive hematopoietic stem and progenitor cells(33). This included marked upregulation of *RUNX1*, which is essential for definitive hematopoietic development(3, 21, 32). Additional analysis of cell fate potential by further differentiation revealed the multilineage potential of our hematopoietic progenitors. We have demonstrated that *SOX17* forward programming results in an EHT and activation of hematopoietic gene networks.

In summary, this is the first report of forward programming of hPSCs to HE cells with SOX17. To our knowledge this is also the first evidence of a single TF mediating an EHT with human cells in the absence of small molecules and growth factors. Furthermore, these findings place SOX17 in a place of heightened importance in human hematopoietic development and demand further investigation. For example, additional study of triphasic SOX17 expression control, mimicking existing evidence from murine development, could lead to further improvements for *in vitro* forward programming and new insights for human HSC emergence(39). We expect these findings will increase our understanding of human hematopoietic development and lead to improved manufacturing of therapeutically relevant cells.

## Materials and Methods

### Maintenance of hPSCs

Human embryonic stem cells (H9, SOX17-mCherry H9 (21), OCT4-GFP H1) and induced pluripotent stem cells (6-9-9) were maintained on either Matrigel (Corning) or iMatrix-511 silk (Nacalai USA) coated plates in LaSR or mTeSR1 (Stemcell Technologies) pluripotent stem cell medium according to previously published methods (12, 40). H9+XLone-SOX17 cells were maintained with 20 µg/mL blasticidin (Sigma) to prevent construct silencing. All drugs were removed upon initiating differentiation or forward programming. Cells were routinely tested to ensure mycoplasma free culture conditions using an established PCR based detection method (41). Cell line details are included in **Table S2**.

### Endothelial progenitor differentiation of hPSCs

Endothelial progenitor differentiation of hPSCs was initiated when hPSCs seeded on Matrigel or iMatrix-511 silk coated plates reached 60% confluence in the presence of Y27632 (Cayman Chemical). Differentiation was performed according to previously published methods (1, 12). Briefly, at day 0, cells were treated with 6 µM CHIR99021 (Cayman Chemical) for 48 hours in LaSR Basal medium, which consists of Advanced DMEM/F12, 2.5 mM GlutaMAX, and 60 μg/ml ascorbic acid, with media refreshed after 24 hours. From day 2-5 cells were maintained in LaSR Basal medium with 50 ng/ml VEGF. Analysis was performed on D5. For forward programming experiments, Dox was added from D0-D2 at 1 µg/mL and from D3-D5 at 5 µg/mL. Cells were re-plated on iMatrix-511 coated plates on D2 by dissociation with Accutase (Innovative Cell Technologies) for 5 minutes at 37C, pelleting, and resuspension in the D2 media with 5 µM Y27632.

### Hematopoietic progenitor differentiation of hPSCs

On day 5, cells were re-plated by dissociation with Accutase for 5 minutes at 37C, pelleting, and resuspension in LaSR basal medium with 5 µM Y27632. On D6, media was changed with StemLine II media (Sigma). On D8, the media volume was doubled with fresh media. On D10, the top half of the media was very carefully aspirated and replaced with fresh media. SCF and FLT3L were optional from D6 to D10.

### Single cell RNA sequencing

H9 cells were differentiated using endothelial progenitor differentiation protocol described above. On day 5 of differentiation, cells were treated with Accutase for 10 minutes. Single cells were counted and resuspended in PBS with 0.04% BSA. A cell strainer was used to get rid of debris and clumps of cells. The single cell library was constructed using the Chromium Next GEM Single Cell 3’ protocol. Then the library was sequenced on a NextSeq 550 equipment with the High Output 150 cycle kit. scRNA sequencing data was processed through the 10X Genomics Cell Ranger pipeline to generate count matrices. These count matrices were then analyzed using Seurat version 3.2.1 (42, 43). Briefly, quality control filters were applied to sort out dying or dead cells and multiplets. The gene expression for each cell was then normalized by the total expression, scaled and log transformed. A linear transform was then applied to the data prior to dimensional reduction via PCA analysis. Statistical (JackStraw) and heuristic (elbow plot) strategies were used to determine the number of principle components to include.

### Immunostaining

Cells were fixed with 4% formaldehyde (Sigma) for 15 min at room temperature. Cells were washed 3 times with PBS and then blocked for 1 hour at room temperature with DPBS with 0.4% Triton X-100 and 5% non-fat dry milk (BioRad). Cells were stained with primary and secondary antibodies (**Table S3**) in DPBS with 0.4% Triton X-100 and 5% non-fat dry milk. Nuclei were stained with Hoechst 33342 (Thermo Fisher Scientific). A Nikon TI Eclipse epifluorescence microscope was used for image capture and analysis. Fiji and Matlab were used for further analysis and quantification.

### Flow cytometry analysis

For staining and analysis of fixed cells, after dissociation with TrypLE Express (differentiated cells) or Accutase (hPSCs), cells were pelleted and resuspended in DPBS with 1% formaldehyde for 30 minutes at room temperature. Cells were pelleted and washed 3 times with DPBS. Cells were stained with primary and secondary antibodies (**Table S3**) in DPBS with 0.1% Triton X-100 and 0.5% BSA for 2 hours at room temperature. Then cells were pelleted and washed 3 times with DPBS with 0.5% BSA before analysis.

For staining and analysis of live cells, cells were dissociated with TrypLE Express (differentiated cells) or Accutase (hPSCs) and pelleted. For suspension cultures of hematopoietic progenitors, cells were filtered with a 100 µm cell strainer and pelleted. Cells were then resuspended in DPBS with 0.5% BSA and the appropriate conjugated primary antibody dilution and incubated at room temperature for 30 minutes. Cells were pelleted and washed with DPBS with 0.5% BSA. Data were collected on a BD Accuri C6 Plus flow cytometer and analyzed using FlowJo. Gating was based on the corresponding untreated or secondary antibody stained cell control.

### Western blotting

Cells were washed with DPBS and lysed with Mammalian Protein Extraction Reagent (Thermo Fisher) with 1X Halt’s Protease and Phosphatase (Thermo Fisher) by incubation for 3 minutes. Cell lysate was collected and stored at -80 C until used. Samples were mixed with Laemmli sample buffer (BioRad) at a working concentration of 1X and incubated at 97 °C for 5 minutes. Samples were loaded into a pre-cast MP TGX stain free gel (BioRad) and run at 200V for 30 minutes in 1X Tris/Glycine/SDS buffer (BioRad). Protein was transferred to a PVDF membrane using a Trans-blot Turbo Transfer System (BioRad). The membrane was blocked for 30 minutes at room temperature in 1X TBST with 5% Dry Milk. The membrane was incubated overnight at 4C with primary antibodies and for 1 hour at room temperature with secondary antibodies (**Table S3**) in 1X TBST with 5% Dry Milk. The membrane was washed between each antibody exposure with 1X TBST. Chemiluminescence was activated using Clarity Western ECL Substrate (BioRad) and the blot was imaged using a ChemiDoc Touch Imaging System and Image Lab software (BioRad). Blots were analyzed using Fiji software.

### Quantitative PCR (qPCR)

RNA was extracted from cells using a Direct-zol RNA MiniPrep Plus Kit (Zymo Research R2071). A Maxima First Strand cDNA Synthesis kit (Thermo Fisher K1641) was used to generate cDNA. A BioRad CFX Connect system was used for performing qPCR with PowerSYBR Green PCR Master Mix (Applied Biosystems 4367659) and primers (**Table S4**). Data was analyzed by the ΔΔCt method where target Ct values were normalized to GAPDH Ct values and fold changes in target gene expression were determined by comparing to day 0 samples. Each sample was run in triplicate. In the event that no measurable expression was detected, relative expression to GAPDH was set to zero.

### Generation of H9 XLone-SOXF cells

The open reading frame for human SOX17 was PCR amplified using GoTaq Master Mix (Promega) from the PB-TRE3G-SOX17 plasmid (**Table S5**). The amplicon was gel purified and ligated into XLone, which was linearized using restriction enzymes KpnI and SpeI (New England Biolabs), using In-Fusion ligase (TaKaRa Bio). XLone-SOX7 and XLone-SOX18 were cloned into XLone by Genewiz. To generate transgenic cell lines, hPSCs were dissociated with Accutase for 10 minutes at 37C and pelleted. The cell pellet was resuspended in 100 µL PBS with 8 µg of plasmid DNA, including 3 µg EF1a-hyPBase and 5 µg XLone-SOX17 (**Table S5**). The mixture was transferred to a cuvette and nucleofected using the CB150 program on the Lonza 4D Nucleofector. All plasmid DNA used was prepared using an Invitrogen PureLink HiPure Plasmid Filter Midiprep Kit. Cells were plated at a high density with 5 µM Y27632. Successfully modified cells were purified using media supplemented with 30 µg/mL blasticidin. Upon achieving a relatively pure population, cells were maintained in media containing 20 µg/mL blasticidin. All plasmids generated have been submitted to Addgene.

### Hematopoietic Colony Forming Unit Assay

5 ×10 3 Floating cells collected at different time points (day 8, day 10, and day 12) were grown in 1 mL of cytokine containing MethoCult H4434 medium (StemCell Technologies, Vancouver) at 37° C. After 14 days, the hematopoietic colonies were scored for colony-forming units (CFUs) according to cellular morphology.

### Giemsa Staining

Day 10 floating cells were collected and methanol fixed on a glass slide. The slide was then stained for 60 minutes at room temperature in a 1:20 dilution of Giemsa stain solution (Sigma-Aldrich). Cells were then washed and mounted for imaging.

### Statistics

Experiments were performed in triplicate. Data obtained from multiple experiments or replicates are shown as the mean ± standard error of the mean. Where appropriate, one or two tailed Student’s *t* test or ANOVA was utilized (alpha = 0.05) with a Bonferroni or Tukey’s post hoc test where appropriate. Data were considered significant when p < 0.05. Statistical tests were performed using MATLAB or GraphPad Prism.

## Acknowledgments

The authors would like to acknowledge and thank the Penn State Genomics Core Facility staff for their support and assistance with completing the scRNA sequence experiments. We also thank Dr. Pentao Liu’s lab for kindly sharing their EF1a-hyPBase plasmid. This work was supported by NIH Trailblazer Award R21EB026035 to X.L.L. and NSF CAREER Award 1943696 to X.L.L. The data sets obtained and used in this study are available upon request submitted to the corresponding author. High-throughput sequencing data obtained in this study has been submitted to GEO and is available under the accession number GSE161408.

## Author Contributions

L.N.R performed experiments, analyzed data, and wrote the manuscript. Y.J., Y.C., and X.B. performed experiments and analyzed data. X.L. performed experiments, analyzed data, wrote the manuscript and supervised the project.

