## Supplemental Information for "Direct induction of hemogenic endothelial progenitors from hPSCs by defined factors revealed by single-cell transcriptome analysis" (from another version of this preprint: version 1, 2021-03-19)

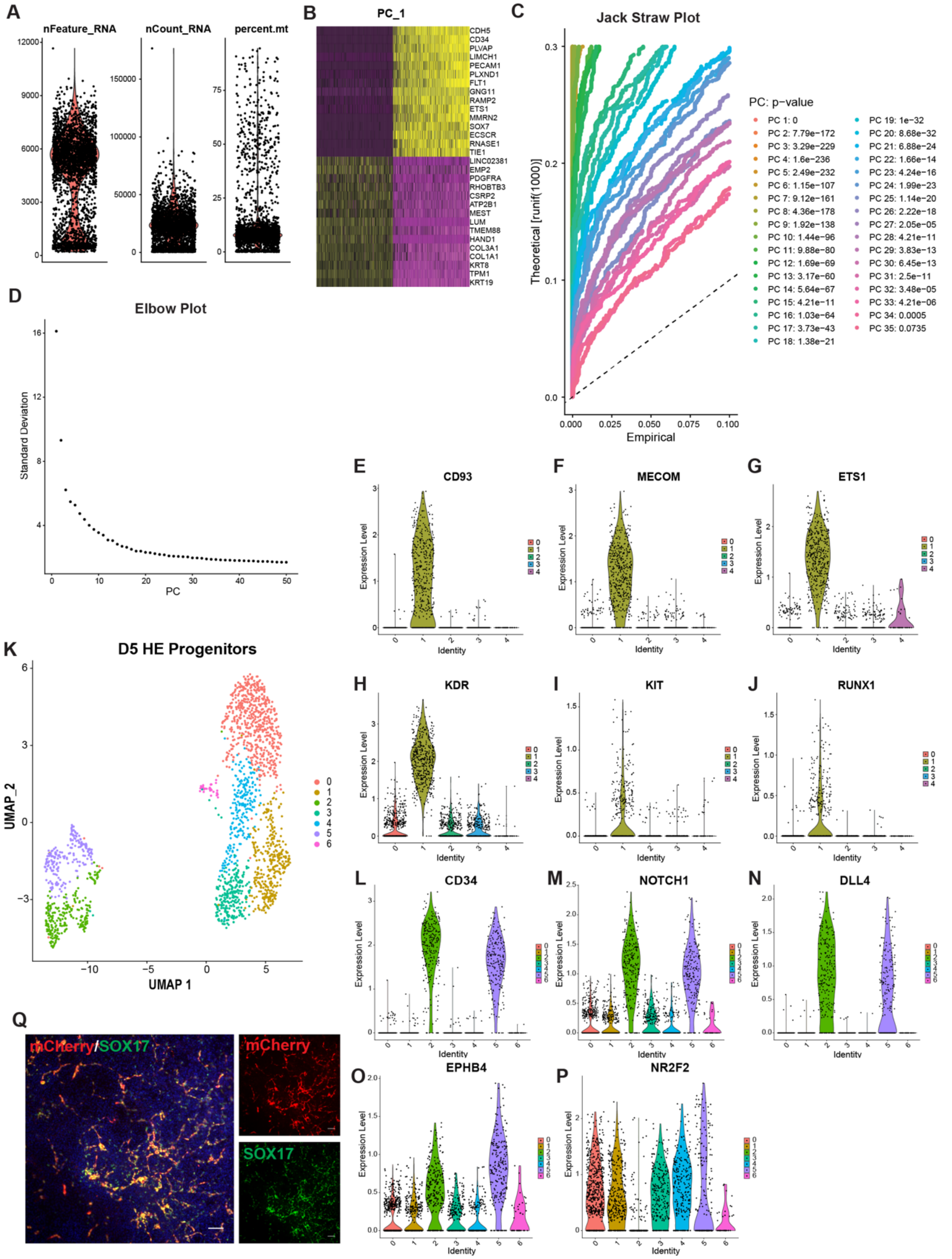


**Fig. S1. Single cell RNA sequencing data analysis and validation. A)** Violin plots indicating the number of genes, counts, and percentage of the reads mapped to mitochondrial genes. **B)** Heat map for principal component 1. **C)** Jack Straw plot for the first 50 principal components. **D)** Elbow plot for the first 50 principal components. **E-J)** Violin plots showing expression of HE cell associated genes CD93 (E), MECOM (F), ETS1 (G), KDR (H), KIT (I), and RUNX1 (J). **K)** UMAP projection with increased resolution subdividing cluster 1 into two (clusters 2 and 5). **L-Q)** Violin plots showing expression of HE and endothelial type associated genes CD34 (L), NOTCH1 (M), DLL4 (N), EPHB4 (O), NR2F2 (P). **Q) I**mmunofluorescence showing mCherry reporter accuracy by comparison with antibody staining of SOX17. Scale bars are 100 μm.

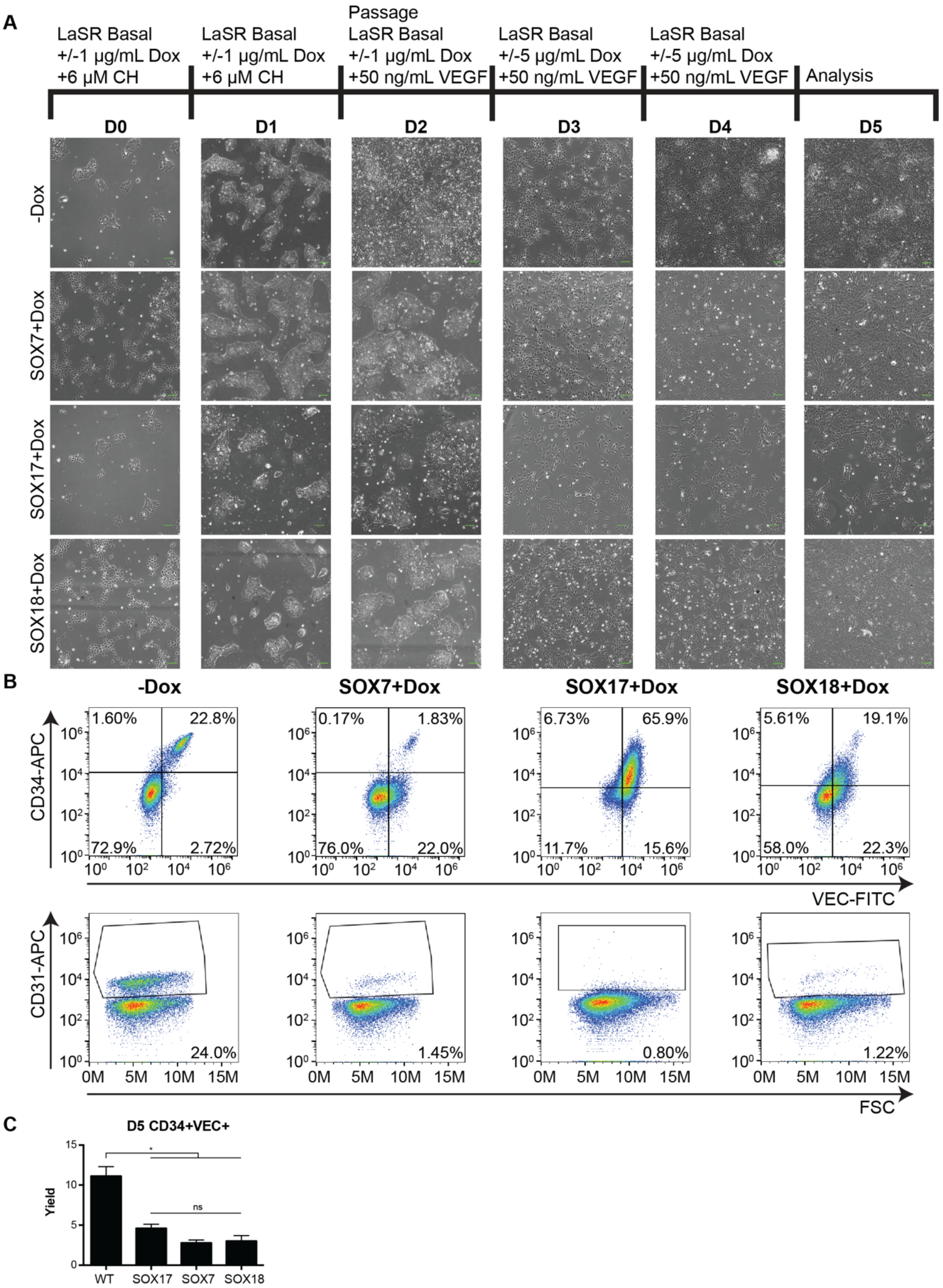


**Fig. S2. Day 5 cells resulting from differentiation with SOXF overexpression. A)** Brightfield images of control, H9+XLone-SOX7, H9+XLone-SOX17, and H9+XLone-SOX18 cells differentiated to day 5 HE cells with and without Dox. Scale bars are 100 µm. **B)** Representative flow cytometry plots analyzing CD34, VEC, and CD31 expression in day 5 cells (n=3). **C)** Yield plot for day 5 cells. Error bars represent standard error of the mean. * indicates $p < 0.05$.

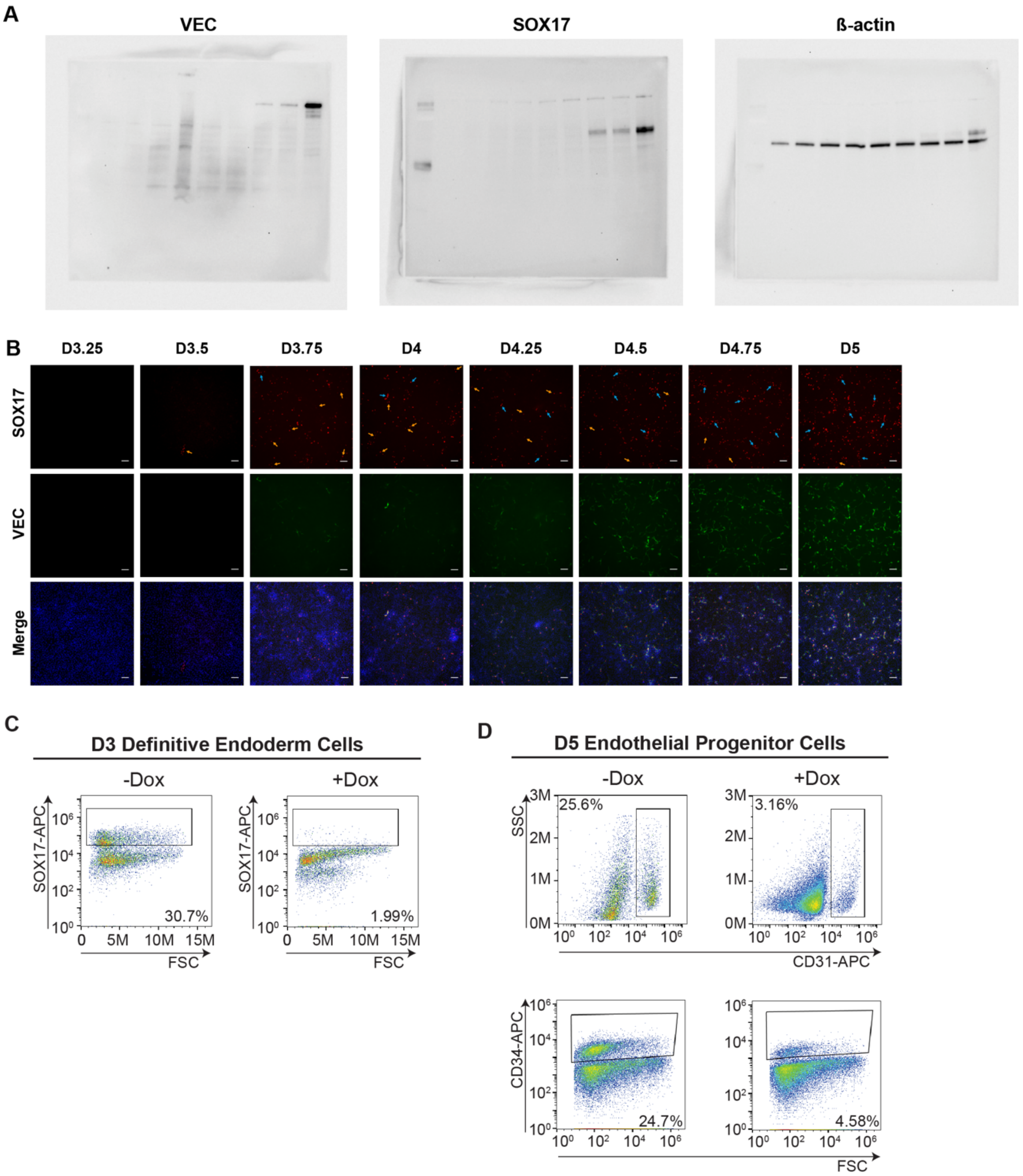


**Fig. S3. SOX17 expression occurs prior to VEC expression and repression prevents HE differentiation. A)** Images showing full blots for western blots shown in Figure 3. **B)** Representative immunofluorescent images of cells every six hours of the differentiation. Orange arrows indicate SOX17$^+$ cells and blue arrows indicate SOX17$^+$VEC$^+$ cells. Scale bars are 100 µm. **C)** Flow analysis of D3 definitive endodermal cells differentiated with and without Dox treatment. **D)** Representative flow analysis plots showing CD31 and CD34 expression in D5 endothelial progenitors differentiated with and without Dox treatment (n=3).

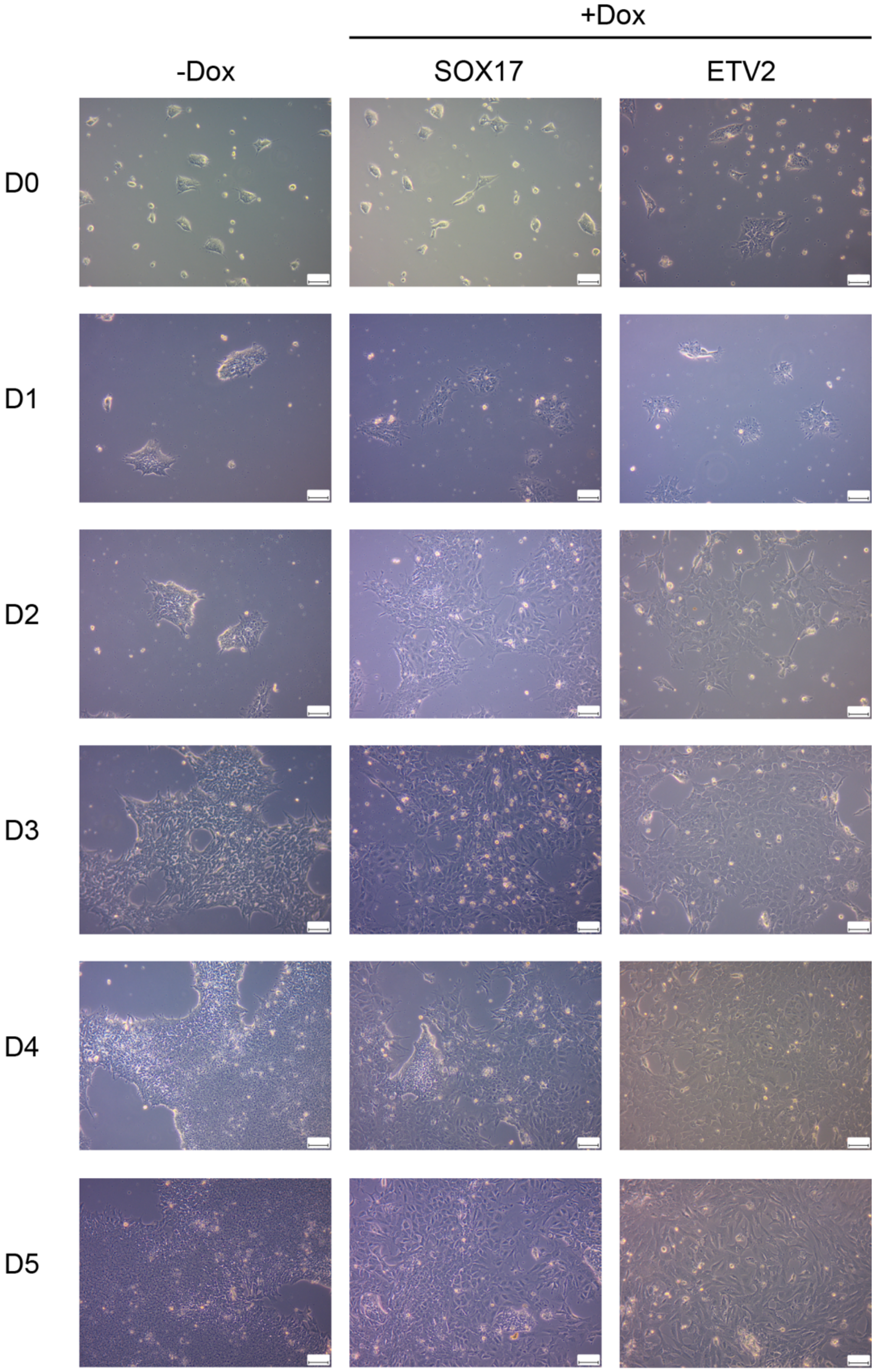

**Fig. S4. SOX17 and ETV2 forward programming.** Daily brightfield images showing cells treated without Dox or with Dox for SOX17 and ETV2 overexpression cell lines. Scale bars are 130 µm.

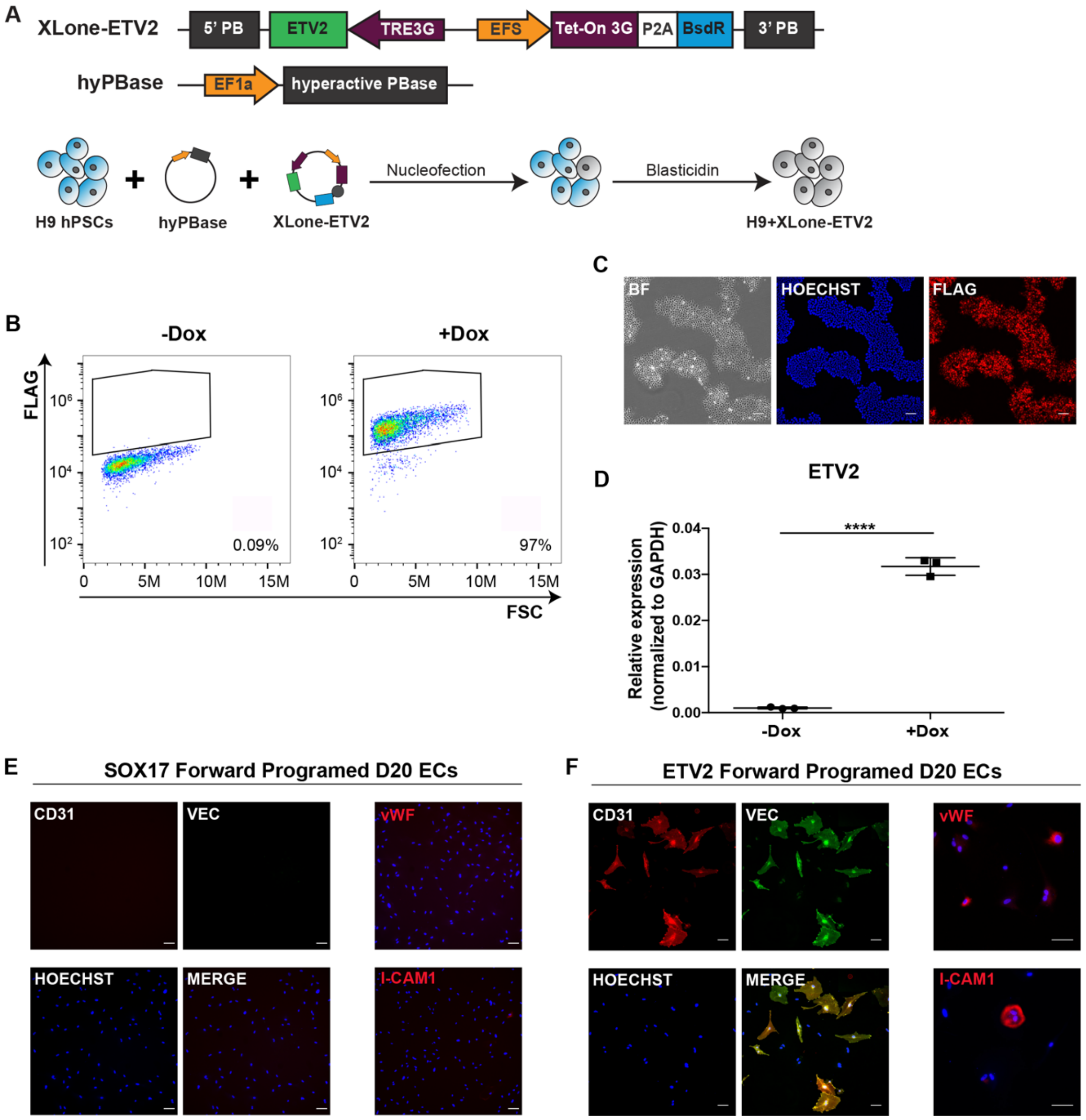


**Fig. S5. Generation of hPSCs with inducible ETV2 overexpression and comparison of SOX17 and ETV2 forward programming mature endothelial cell potential. A)** Schematic showing the generation of H9+XLone-ETV2 cells. **B)** Flow analysis showing ETV2 associated FLAG expression with and without 24 hours of Dox treatment. **C)** Immunofluorescence showing ubiquitous FLAG expression associated with ETV2. Scale bars are 100 μm. **D)** qPCR data confirming an increase in ETV2 expression with Dox treatment (n=3). **** indicates $p < 0.0001$. Error bar represents standard error of the mean. **E and F)** Immunofluorescence analysis of D5 SOX17 (E) and ETV2 (F) forward programmed cells that were further differentiated to D20 in commercially available endothelial cell differentiation media. Scale bars are 100 μm.

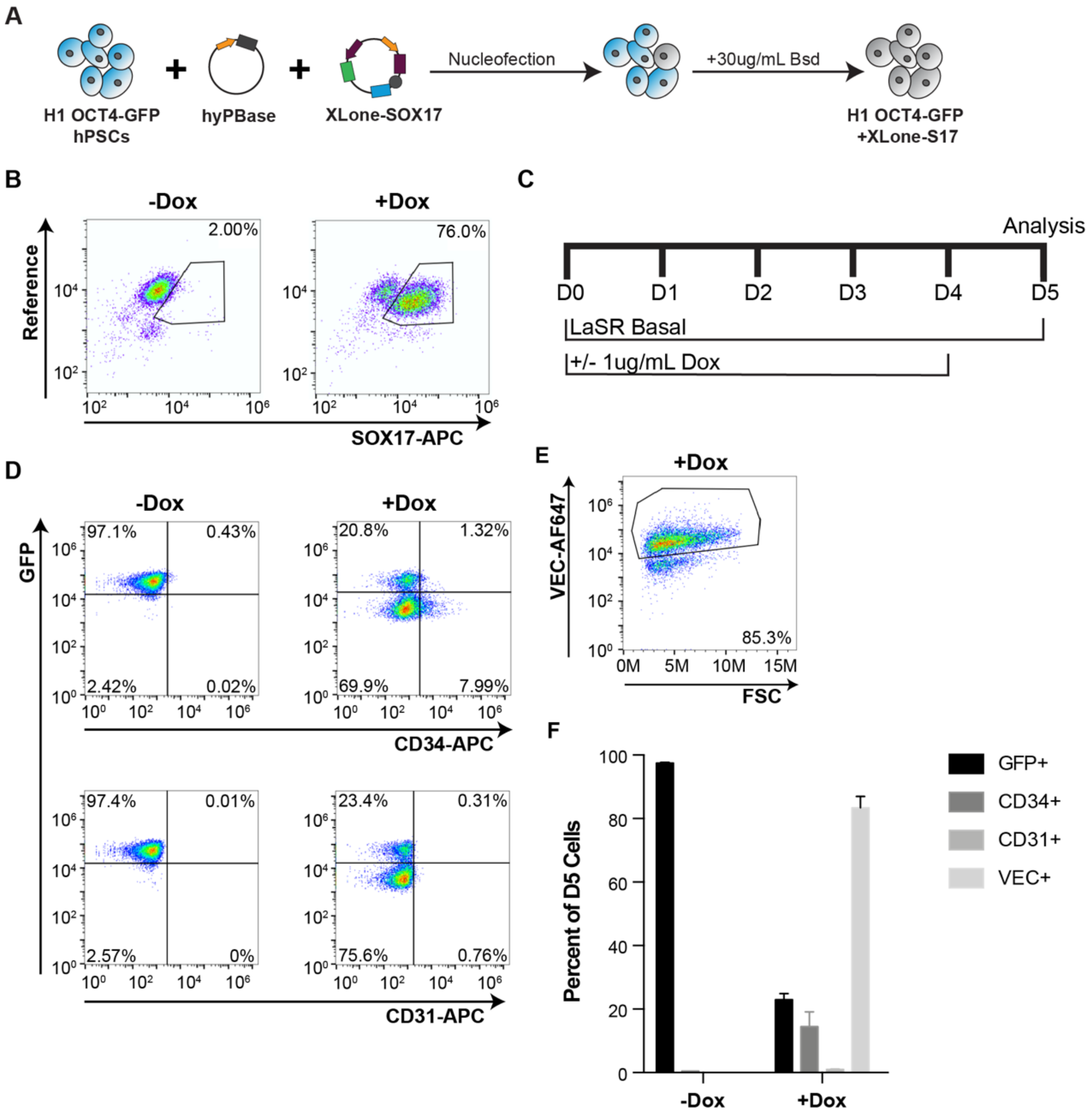


**Fig. S6. H1+XLone-SOX17 forward programming. A)** Schematic showing the generation of H1 OCT4-GFP+XLone-SOX17 cells. **B)** Flow analysis showing SOX17 expression with and without 24 hours of Dox treatment. **C)** Experimental schematic of forward programming protocol. **D and E)** Representative flow analysis of CD34, CD31, GFP (D), and VEC (E) expression for day 5 forward programmed cells (n=3). **F)** Quantification of flow cytometry data (n=3). Error bars represent standard error of the mean.

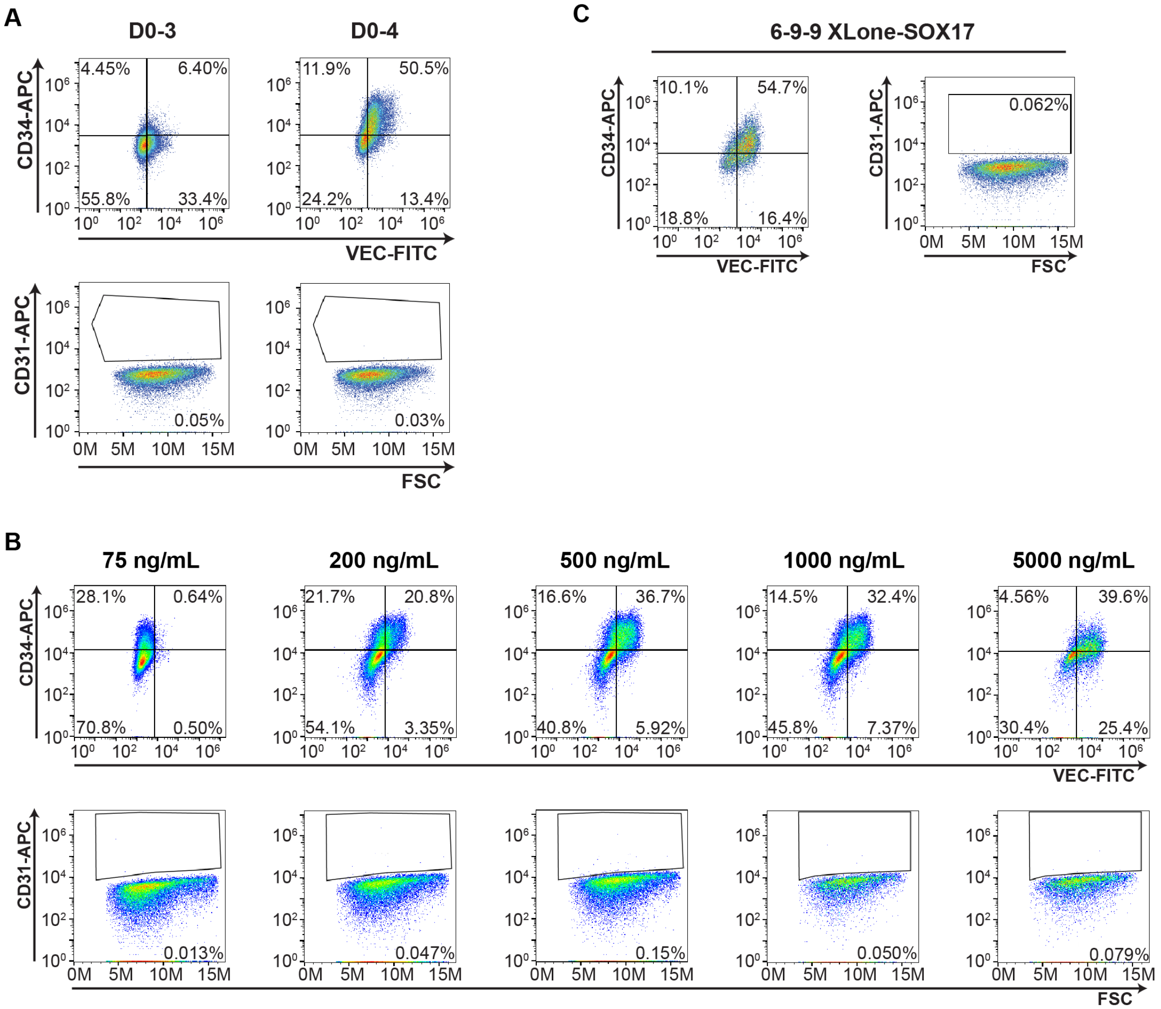


**Fig. S7. SOX17 forward programming temporal and concentration optimization. A)** Representative flow cytometry analysis for CD34, CD31, and VEC of SOX17 d5 forward programmed cells treated with Dox for 3 or 4 days (n=3). **B)** Representative flow cytometry analysis of CD34, CD31, and VEC for SOX17 d5 forward programmed cells treated with varied concentrations of Dox for 5 days (n=3). **C)** Flow cytometry analysis for CD34, CD31, and VEC of 6-9-9+XLone-SOX17 day 5 forward programmed cells treated with Dox for 5 days and passaged on day 2.

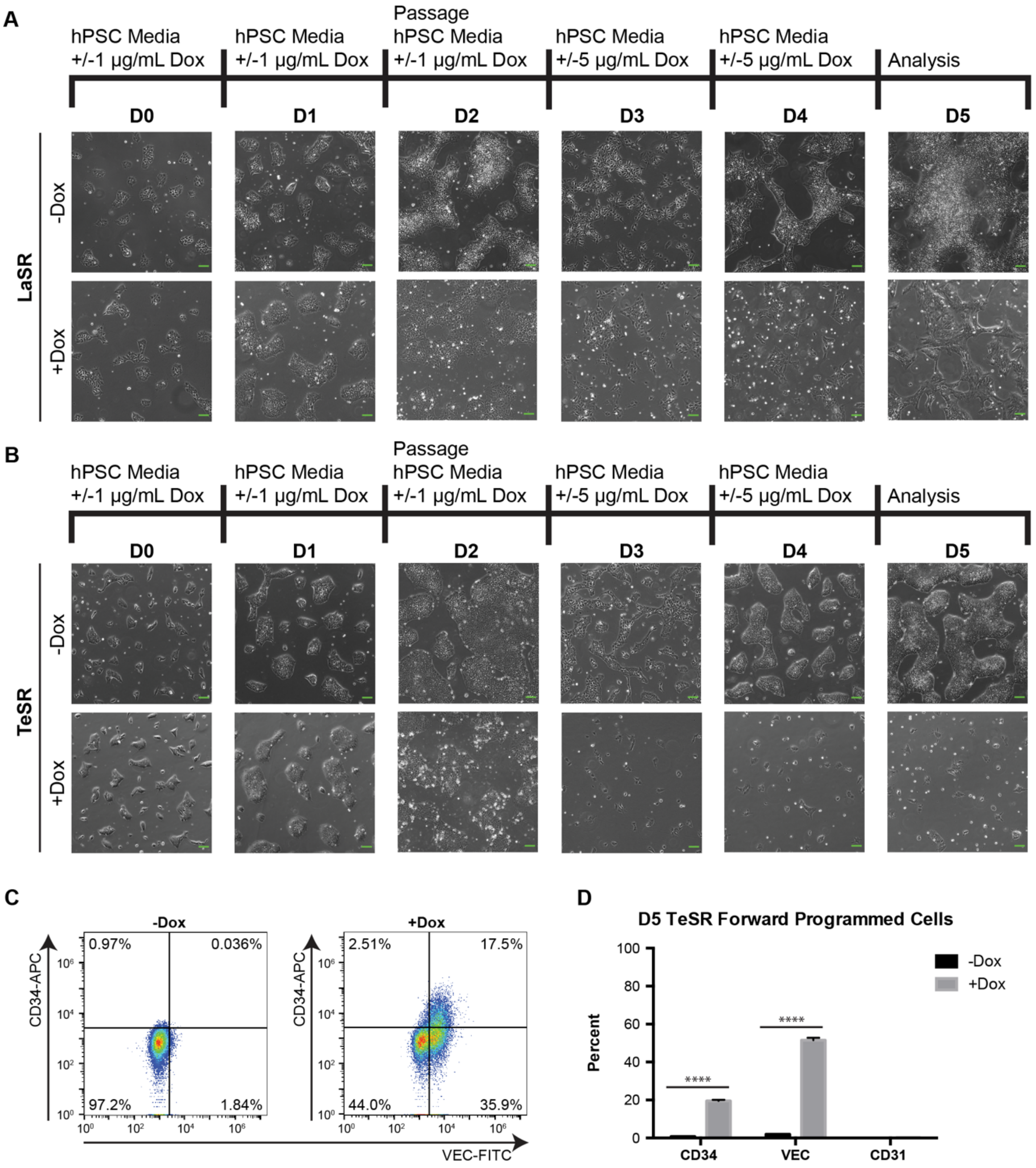


**Fig. S8. SOX17 forward programming in hPSC media. A)** Bright field images of cells forward programmed with or without SOX17 overexpression in LaSR hPSC media. Scale bars are 100 μm. **B)** Bright field images of cells forward programmed with or without SOX17 overexpression in mTeSR1 hPSC media. Scale bars are 100 μm. **C)** Representative flow cytometry analysis of CD34 and VEC expression in day 5 cells (n=3). **D)** Quantification of flow cytometry analysis for CD34, CD31, and VEC in day 5 cells (n=3). **** indicates $p < 0.0001$. Error bars represent standard error of the mean.

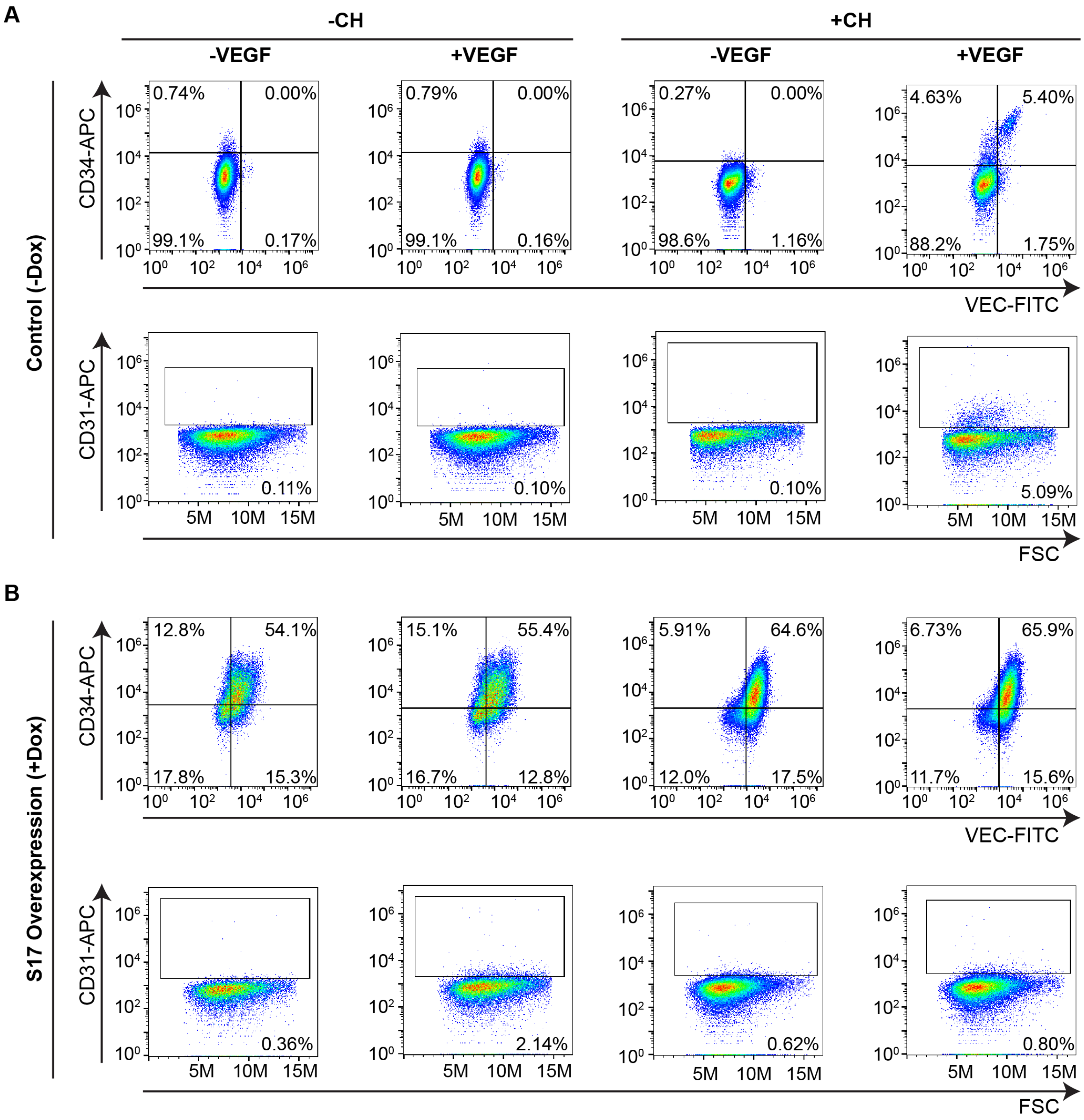


**Fig. S9. Combinatorial testing of SOX17 forward programming and HE differentiation factors for HE specification.** Representative flow cytometry plots for day 5 cells differentiated with stepwise inclusion of HE specification factors without (A) or with (B) Dox treatment (n=3).

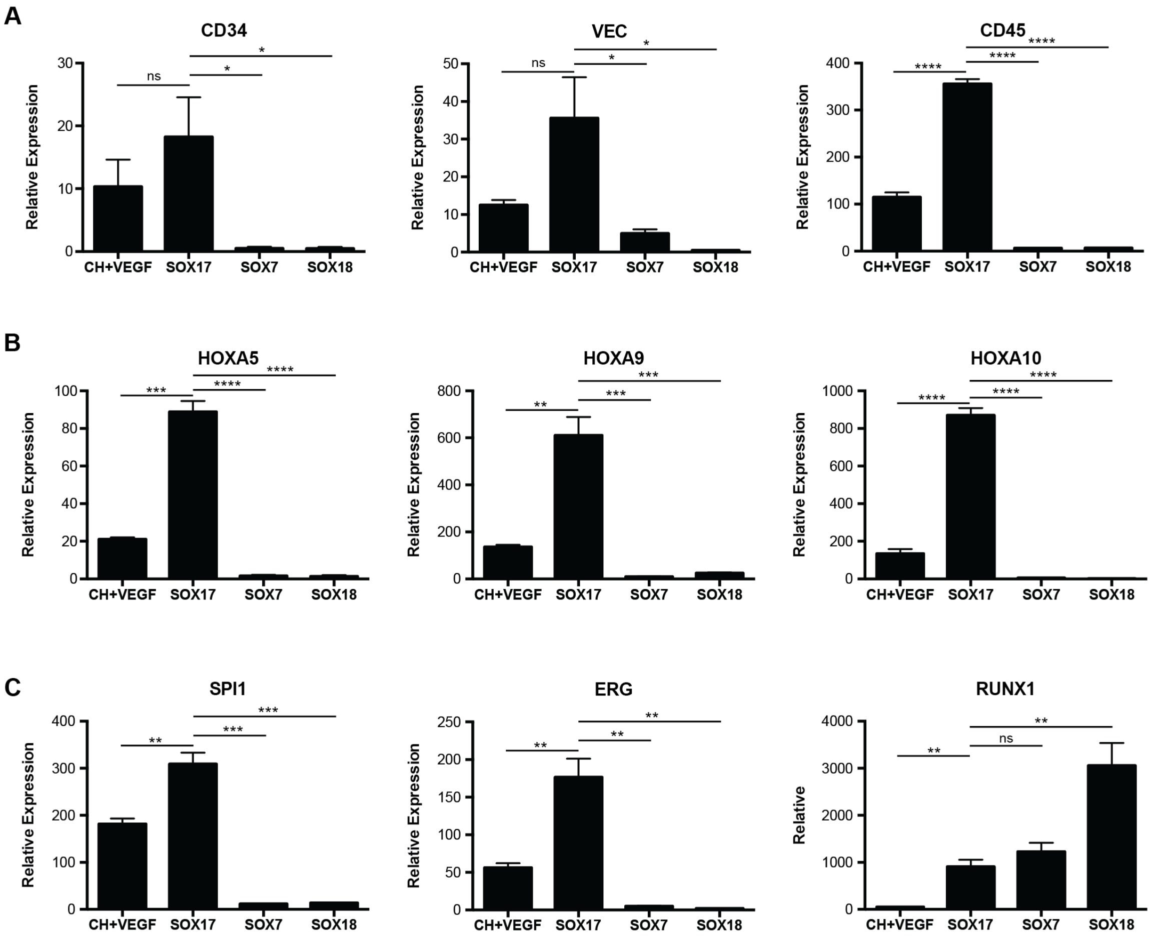


**Fig. S10. Comparative analysis of hematopoietic factors in differentiated and SOXF forward programmed cells.** Quantification of gene expression determined by qPCR (n=3) for CD34 (A), VEC (A), CD45 (A), HOXA5 (B), HOXA9 (B), and HOXA10 (B), SPI1 (C), ERG (C), RUNX1 (C). *, **, ***, and **** indicates $p < 0.05$, 0.01, 0.001, and 0.0001 respectively. Error bars represent standard error of the mean.

**Table S2.** Cell Lines

| Cell Line Name | Source | Sex | Cell Type |
|---|---|---|---|
| H9 | WiCell | F | hESC |
| 6-9-9 | WiCell | M | iPSC |
| H9 SOX17-mCherry | These cells were a generous gift from Ed Stanley. | F | Genetically modified knock in reporter hESC |
| H1 OCT4-GFP | WiCell | M | Genetically modified knock in reporter hESC |

**Table S3.** Antibodies

| Antibody | Source/Host and Isotype/Clone/Catalog# | Application |
|---|---|---|
| SOX17-APC | R&D Systems/ goat IgG/ IC1924A | 1:50 (IF/FC) |
| VE-Cadherin | Santa Cruz/ mouse $IgG_1$/ Clone: F-8/ sc-9989 | 1:100 (IF/FC)<br>1:1000 (WB) |
| CD144-FITC | MACS/ recombinant human $IgG_1$/ Clone REA199/ 130-100-742 | 1:100 (FC) |
| CD34-FITC | MACS/ mouse IgG2a / Clone AC136/ 130-113-178 | 1:100 (FC) |
| CD34-APC | MACS/ mouse IgG2a / Clone AC136 / 130-113-176 | 1:100 (FC) |
| SOX17 | R&D Systems/ rabbit IgG/ IC1924A | 1:20000 (WB) |
| β-actin-HRP | Cell Signaling Technology/ rabbit/ Clone: 13E5/ 5125S | 1:20000 (WB) |
| CD31-APC | MACS/ mouse / Clone:AC128 /130-092-652 | 1:100 (IF/FC) |
| CD73-PE | Biolegend/ mouse IgG1 / Clone:AD2 /344003 | 1:100 (FC) |
| vWF | Santa Cruz/ mouse IgG1 / Clone:F8/86 / sc-53466 | 1:200 (IF) |
| I-CAM1 | DSHB/ mouse IgG1 / p2a4 | 1:300 (IF) |
| FLAG | Sigma-Aldrich/ rabbit IgG / F7425 | 1:500 (FC) |
| CD45-APC | BIolegend/ Clone 2D1 / 368511 | 1:20 (FC) |
| CD44-FITC | Biolegend/ Clone BJ18 / 338803 | 1:20 (FC) |
| Secondary Antibody | Alexa 488 Goat anti Ms IgG/ A-11029 | 1:1000 |
| Secondary Antibody | Anti-mouse IgG HRP-linked/ CST 7076S | 1:1000 |
| Secondary Antibody | Goat IgG HRP-conjugated/ R&D Systems HAF017 | 1:1000 |
| Secondary Antibody | Alexa 647 Goat anti Rb IgG/ A-21244 | 1:1000 |
| Secondary Antibody | Alexa 647 Goat anti Ms IgG/ A-21235 | 1:1000 |

**Table S4.** qPCR primers

| Primer | Sequence |
|---|---|
| GAPDH_FWD | 5’-GTGGACCTGACCTGCCGTCT-3’ |
| GAPDH_REV | 5’-GGAGGAGTGGGTGTCGCTGT-3’ |
| OCT4_FWD | 5’-CAGTGCCCGAAACCCACAC-3’ |
| OCT4_REV | 5’-GGAGACCCAGCAGCCTCAAA-3’ |
| SOX2_FWD | 5’-CAAGATGCACAACTCGGAGA-3’ |
| SOX2_REV | 5’-GTTCATGTGCGCGTAACTGT-3’ |
| SOX17_FWD | 5’-GGCGCAGCAGAATCCAGA-3’ |
| SOX17_REV | 5’-CCACGACTTGCCCAGCAT-3’ |
| GATA2_FWD | 5’-CTCCCAGCTCTACTCCAGG-3’ |
| GATA2_REV | 5’-GTGGTGTGAGTCGGGGTG-3’ |
| CD31_FWD | 5’-GCTGACCCTTCTGCTCTGTT-3’ |
| CD31_REV | 5’-TGAGAGGTGGTGCTGACATC-3’ |
| VEC_FWD | 5’-GTGTTCACGCATCGGTTGTT-3’ |
| VEC_REV | 5’-CAAATGTGTACTTGGTCTGGGTG-3’ |
| CD34_FWD | 5’-CCTAAGTGACATCAAGGCAGAA-3’ |
| CD34_REV | 5’-GCAAGGAGCAGGGAGCATA-3’ |
| DLL4_FWD | 5’-GCACTCCCTGGCAATGTACT-3’ |
| DLL4_REV | 5’-GCCATCCTCCTGGTCCTTAC-3’ |
| TAL1_FWD | 5’-GGGATCTGATTCTATCGCCCA-3’ |
| TAL1_REV | 5’-GCTTTCCCCCTTTTTCGCTG-3’ |
| SPI1_FWD | 5’-GCCCTGCAATGTCAAGGGA-3’ |
| SPI1_REV | 5’-GAAGTCCCAGTAATGGTCGCT-3’ |
| ETS1_FWD | 5’-CATCCACAAGACAGCGGGG-3’ |
| ETS1_REV | 5’-CTCGTCGGCATCTGGCTTG-3’ |
| RUNX1_FWD | 5’-GGTTTCGCAGCGTGGTAAAA-3’ |
| RUNX1_REV | 5’-GCACTGTGGGTACGAAGGAA-3’ |
| CD43_FWD | 5’-CATTCCTCAGCCAAGAGCCA-3’ |
| CD43_REV | 5’-GCAAGAGCTGGGACCTGAG-3’ |
| CD45_FWD | 5’-CAACAGTGGAGAAAGGACGC-3’ |
| CD45_REV | 5’-TGTCCAGAAAGGCAAAGCCA-3’ |
| LCOR_FWD | 5’-CAGCAATGCTCCGTTGAGAG-3’ |
| LCOR_REV | 5’-GAGGGAAACGGACCTACAC-3’ |
| ERG_FWD | 5’-CTTGGTCGGAATGGGGAGAG-3’ |
| ERG_REV | 5’-TGTTCAGAACCTGACGGCTTT-3’ |
| HOXA5_FWD | 5’-GCTGCACATAAGTCATGACAACA-3’ |
| HOXA5_REV | 5’-CGCTCAGATACTCAGGGACG-3’ |
| HOXA9_FWD | 5’-CCCATCGATCCCAATAACCC-3’ |
| HOXA9_REV | 5’-TGTGGCCTGAGGTTTAGAGC-3’ |
| HOXA10_FWD | 5’-TACTTCCGCCTTTCTCAGGC-3’ |
| HOXA10_REV | 5’-CCTTTGGAATTGCCCAGGGA-3’ |

**Table S5.** Plasmids used

| Plasmid | Source | Addgene # |
| --- | --- | --- |
| XLone | This plasmid was generated in the Lian Lab. | 96930 |
| XLone-SOX17 | This plasmid was generated in the Lian Lab. | Pending |
| XLone-SOX7 | This plasmid was generated in the Lian Lab. | Pending |
| XLone-SOX18 | This plasmid was generated in the Lian Lab. | Pending |
| XLone-ETV2 | This plasmid was generated in the Lian Lab. | Pending |
| XLone-Cas13d-SOX17-gRNA | This plasmid was generated in the Lian Lab. | Pending |
| PB-TRE3G-SOX17 | This plasmid was a gift from David Vereide. | 104541 |
| EF1a-hyPBase | This plasmid was a gift from Pentao Liu. | NA |
